# Sweeps in space: leveraging geographic data to identify beneficial alleles in *Anopheles gambiae*

**DOI:** 10.1101/2025.02.07.637123

**Authors:** Clara T. Rehmann, Scott T. Small, Peter L. Ralph, Andrew D. Kern

**Affiliations:** University of Oregon, Institute of Ecology and Evolution; University of Oregon, Department of Biology; University of Oregon, Department of Data Science

## Abstract

As organisms adapt to environmental changes, natural selection modifies the frequency of non-neutral alleles. For beneficial mutations, the outcome of this process may be a selective sweep, in which an allele rapidly increases in frequency and perhaps reaches fixation within a population. Selective sweeps have well-studied effects on patterns of local genetic variation in panmictic populations, but much less is known about the dynamics of sweeps in continuous space. In particular, because limited movement across a landscape leads to unique patterns of population structure, spatial dynamics may influence the trajectory of selected mutations. Here, we use forward-in-time, individual-based simulations in continuous space to study the impact of space on beneficial mutations as they sweep through a population. In particular, we show that selection changes the joint distribution of allele frequency and geographic range occupied by a focal allele and demonstrate that this signal can be used to identify selective sweeps. We then leverage this signal to identify in-progress selective sweeps within the malaria vector *Anopheles gambiae*, a species under strong selection pressure from vector control measures. By considering space, we identify multiple previously undescribed variants with potential phenotypic consequences, including mutations impacting known IR-associated genes and altering protein structure and properties. Our results demonstrate a novel signal for detecting selection in spatial population genetic data that may have implications for genomic surveillance and understanding geographic patterns of genetic variation.

## Introduction

Positive selection is a fundamental force in population genetics, and selective sweeps are responsible for many well-known examples of adaptation such as changes in melanin production in humans (Williamson et al., 2007), insecticide resistance in *Drosophila* (Schlenke and Begun, 2004), and major architectural differences between teosinte and maize (Clark et al., 2004). The foundational model of a selective sweep describes how a newly-arisen, positively selected variant increases in frequency and eventually fixes (Fisher, 1930, Kojima and Kelleher, 1962). During this process, neutral variants linked to the sweeping variant also increase in frequency, ultimately reducing variation around the site of the sweeping allele (i.e. genetic hitchhiking; Maynard Smith and Haigh (1974), Kojima and Schaffer (1967)). This observation has been the basis of numerous tests for identifying positive selection in genomic data, including those based on summary statistics (Tajima, 1989, Fay and Wu, 2000), explicit likelihood models (Nielsen et al., 2005, DeGiorgio et al., 2016, Chen et al., 2010, DeGiorgio and Szpiech, 2022), and more recently through the use of supervised machine learning (Schrider and Kern, 2016, Kern and Schrider, 2018, Sugden et al., 2018, Xue et al., 2021, Arnab et al., 2023, Lauterbur et al., 2023). However, this foundational model of a selective sweep - and the methods based on it - assume populations are panmictic and without spatial heterogeniety. While these assumptions simplify models of sweeps, they also overlook potentially confounding variables, as well as those that could better inform analysis. One such factor is that most natural populations live in landscapes where their dispersal and interactions are geographically limited, leading to a near ubiquitous observation in population genetics - isolation by distance (IBD) - that individuals tend to be more closely related the closer they are in geographic space (Wright, 1943). This effect of spatially limited dispersal can have profound impacts on how genetic variation is shared among individuals in a population, both with respect to common summaries of variation such as heterozygosity and the site frequency spectrum, as well as downstream inferences based on that variation such as inferred effective population sizes and genomic association tests (e.g., Battey et al., 2020a). These effects have implications for the ways that evolutionary processes, such as selective sweeps, are identified in natural populations.

Prior investigation into selective sweeps in spatially structured populations has revealed that a beneficial allele will move across space and through the population as a travelling wavefront, historically described in a one-dimensional population by Fisher (1937) and in two dimensions by Kolmogorov et al. (1937). In this model, dispersal distance determines the speed at which a sweep progresses, with smaller dispersal distances slowing the allele’s frequency trajectory relative to the panmictic expectation (Barton et al., 2013). Min et al. (2022) demonstrated how structure across one-dimensional space impacts the population genetic signatures expected after a selective sweep, most notably causing an excess of variants at moderate frequency that were able to recombine onto the sweeping haplotype due to the slower nature of the sweep. Further, Chotai et al. (2024) recently explored the signatures of sweeps in populations structured across continuous two-dimensional space. This study corroborated results from one-dimensional explorations of sweeps in showing how sweeping alleles move more slowly in spatially structured populations and demonstrated that spatial structure “softens” the genetic signals expected from selective sweeps, causing them to appear more like soft sweeps (*c*.*f*., Hermisson and Pennings, 2005). While our understanding of how spatial population structure changes expectations for the genetic signatures left by selective sweeps has increased, particularly in how it may reduce our capacity to detect them in genomic data, it remains unexplored whether this structure can be leveraged to inform genetic analysis of positive selection.

Spatial patterns of genetic variation can reveal information about evolutionary process, rather than simply confound. For example, Battey et al. (2020b) showed how machine learning approaches to geo-referenced data allow for geolocation of genotypes across continuous landscapes, and that the residuals from such predictions can be used to infer historical processes (Battey et al., 2020b, Rehmann et al., 2024). Further, spatial genetic data can be used to infer how organisms move across landscapes (Rousset, 1997, Ringbauer et al., 2017, Al-Asadi et al., 2019, Marcus et al., 2021, Smith et al., 2023, Smith and Kern, 2023) as well as give information about local density of the population (Al-Asadi et al., 2019, Smith et al., 2024). While spatial data has been being utilized to inform genetic analysis of explicitly spatial processes, other fundamental phenomena, such as selective sweeps, are also impacted by spatial population structure. Given that selective sweeps in spatially structured populations behave differently from those in panmictic populations, most notably in their speed and the impact of that on linked neutral variation, this spatial dimension of how genetic variation is shared may contain information that reveals sweeping alleles.

Identifying ongoing selection is of particular importance in populations relevant to human health, such as the malaria-causing parasite *Plasmodium* and its vectors. The malaria vector *Anopheles* is under strong selective pressures from public health interventions (e.g., Bhatt et al., 2015), and thus is frequently reported as having experienced or is currently undergoing selective sweeps in part due to these efforts (Anopheles gambiae 1000 Genomes Consortium, 2017, Clarkson et al., 2021, Xue et al., 2021). Genomic surveillance in these populations is crucial, as variants under positive selection are notably often in genes related to insecticide resistance, (Hemingway et al., 2016, Clarkson et al., 2021), and thus relevant to advances in vector control and prevention of disease transmission. While that is so, the studies identifying these variants often rely on methods that may be confounded by spatial population dynamics, reducing their power to identify sweeping alleles. By using spatial information to inform searches for selective sweeps, we may be able to identify more (or different) variants under positive selection.

In this study, we use forward-in-time simulation to model selective sweeps across a continuous landscape and leverage the joint distribution between an allele’s spatial spread and its frequency to reveal a novel signal of positive selection. We then demonstrate this spatial signal’s potential to reveal alleles under ongoing positive selection both through SNP-centric and windowed approaches, noting that in empirical applications these approaches will identify both globally- and locally-adaptive variants. We apply these methods to a spatially referenced dataset of *Anopheles gambiae* genomes (Ag1000G Selection Atlas, 2017). Our analysis recovers known instances of positive selection in *An. gambiae*, identifying variants within known insecticide-resistant loci, and identifies potentially uncharacterized sources of insecticide resistance as well as other potentially selected variants associated with immune response, sensory perception, and development. Through simulation and empirical analysis, we demonstrate the potential of spatial population structure to identify, rather than obscure, segregating variants under positive selection.

## Results

### A novel spatial signal of selection

How does a beneficial allele move across space? An allele under positive selection that arises at a point along the landscape (position 50 in Figure 1) will increase in frequency where it arises while simultaneously spreading in all directions in the form of a traveling wave (Fisher, 1937, Kolmogorov et al., 1937). While the beneficial allele is globally increasing in frequency, the portion of the landscape occupied by carriers of said allele is constrained by the wavelike dynamics of its spread. In contrast, a neutral allele that arises and is not lost will diffuse in both directions, but generally remain at low or intermediate frequencies across the landscape, as its frequency dynamics are determined by drift (Figure 1). Because positive selection causes a beneficial allele to increase in frequency faster than it can spread through space, these alleles should occupy less space on the landscape than neutral variants at similar global frequencies. We set out to observe this phenomenon in continuous space using forward-in-time, individual-based simulation.

**Figure 1.**
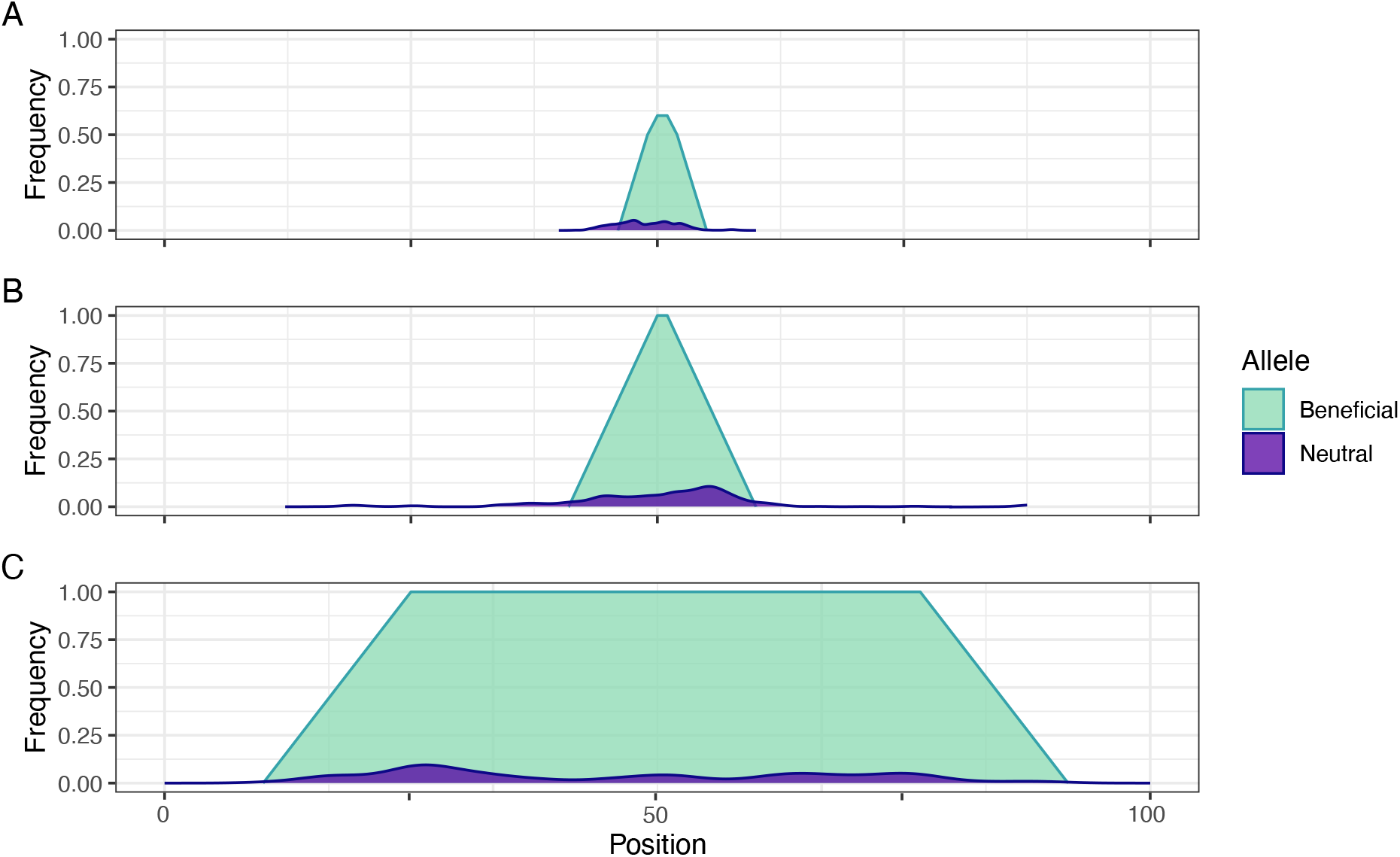
A conceptual model of how positive selection impacts an allele’s spread across a one-dimensional landscape. Panels A, B, and C show a time series of a beneficial allele (in green) and a neutral allele (in blue) arising at position 50 and spreading through space. The frequency of each allele at a given spatial position is indicated along the y axis.

To simulate selective sweeps in continuous space, we used a non-Wright-Fisher framework similar to that described in Chevy et al. (2024), where individuals disperse from their parent across the landscape, compete locally for resources, and reproduce with a spatially proximate mate. Dispersal and interactions are governed by a Gaussian kernel with size *σ*, which represents the average per-generation movement rate, competition distance, and mate search radius (Methods Spatial simulation). On to this continuous space simulation we introduce a positively selected allele – one that reduces individual mortality – to the middle of the simulated genome of an individual at the center of the landscape, then track the progress of the focal mutation until eventual fixation or loss (Figure 2A). Additionally, we compare results of our simulations to simple theoretical deterministic expectations of the spread of a beneficial allele according to traveling wave dynamics (Supplemental section Approximation to Fisher’s wave; Fisher (1937), Kolmogorov et al. (1937)). In our simulations, we tracked the frequency of the allele as well as the area of the landscape occupied by the allele’s carriers, where area was defined by a convex hull formed at the vertices of allele carriers (see Methods).

**Figure 2.**
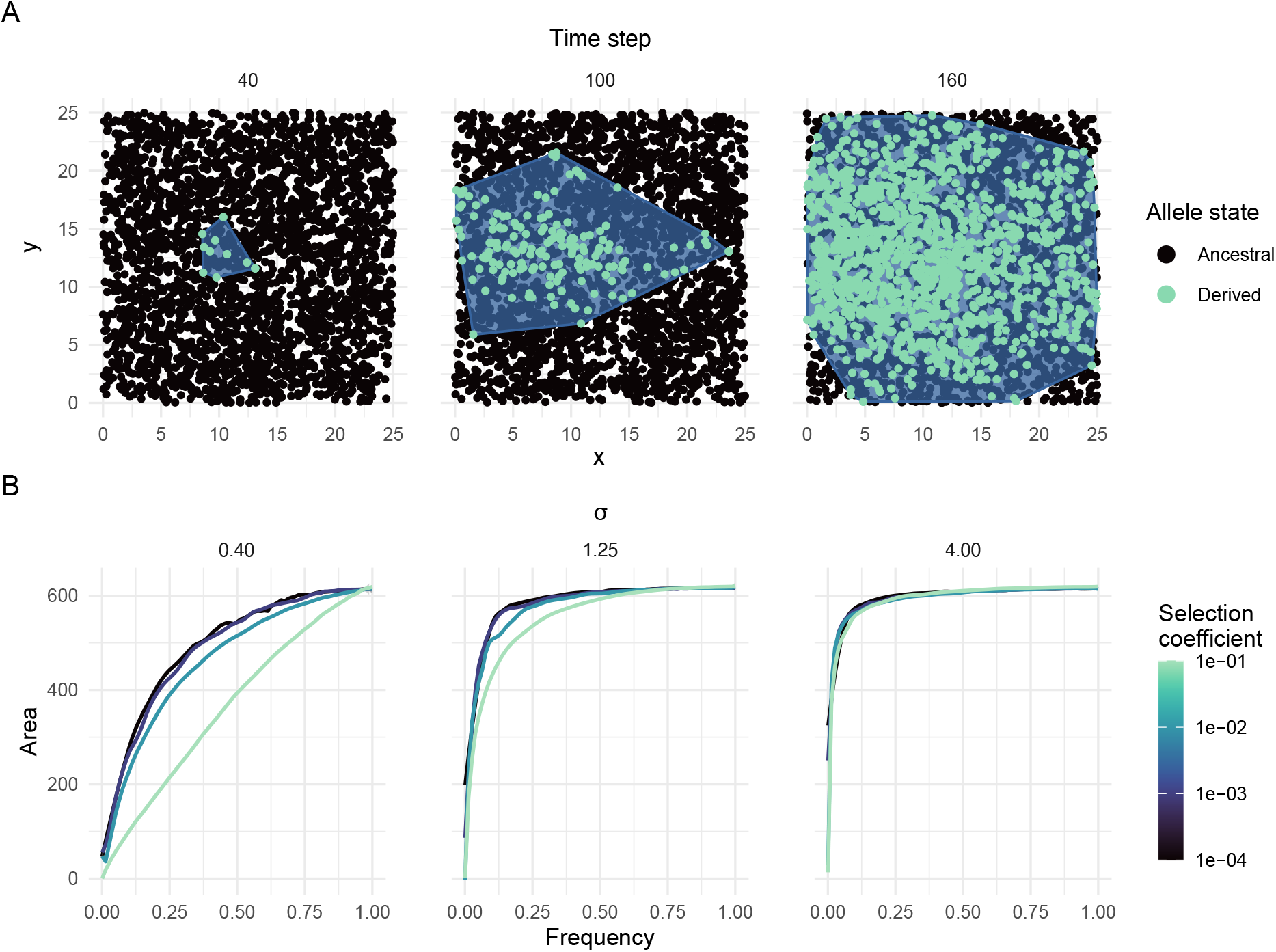
Selective sweeps in continuous space. (A) Example of simulated allele with selection coefficient 0.1 sweeping through a population with a dispersal distance (*σ*) of 0.40. Individuals carrying the sweeping allele are colored in green. (B) Joint frequency-area trajectories of sweeping alleles at dispersal distances 0.40, 1.25, and 4.0. Line color represents the strength of selection; allele area is measured as the area of the polygon encompassing allele carriers.

In Figure 2B we show the allele frequency and area trajectories resulting from our spatial simulations. In accordance with the conceptual model shown in Figure 1, alleles under strong positive selection occupy less space on the landscape in comparison to neutral or weakly selected alleles at similar frequencies. There is a strong relationship between the strength of selection, the frequency of a beneficial allele, and its area: as the strength of selection increases, beneficial alleles cover *less* of the landscape conditional on frequency. This effect is most pronounced at smaller dispersal rates, where the allele’s spread is inherently more limited by spatial population structure, but persists even when per-generation dispersal is high. Selection’s effect on this joint distribution - reducing relative area - is initially counterintuitive as one might expect a sweeping allele to spread more quickly across a landscape than a neutral variant. This is true with respect to time, however conditional on its frequency, a neutral variant will diffuse across the landscape without a substantial increase in frequency. Under positive selection, an allele will both spread across the landscape and increase in frequency, reducing the expected area associated conditional on frequency and creating a visible signal in the joint distribution (Figure 2B). We note that deterministic models of Fisherian wave dynamics recapitulate these dynamics as well (Figure S1), thus drift is not responsible for the general effect.

### Power to detect ongoing selection using space

We next asked whether we could use this spatial signal to identify alleles under ongoing positive selection. Given that positive selection has a striking impact on allele area conditioned on frequency, we decided to use an empirical outlier approach on this joint distribution to localize candidate selected alleles. Simply put, the idea is to identify alleles that occupy less area than expected given their frequency (spatial-frequency or SF outliers). An example of this joint distribution along with a heuristic cutoff to identify SF outliers is shown in Figure 3A. While this distribution is defined based on individual SNPs, sweeping alleles also bring along their linked genetic background, and there may be additional power in aggregating SF outlier signal across windows. To do so, we first identify genome-level SF outliers (as in Fig. 3A) in a genomic window, then define a *z*-score that describes the proportion of variants found within a given window that are SF outliers (see Power analysis section). The goal is to identify particular windows which harbor an excess of spatial-frequency outliers (WSF outliers, see Power analysis; Figure 3B).

**Figure 3.**
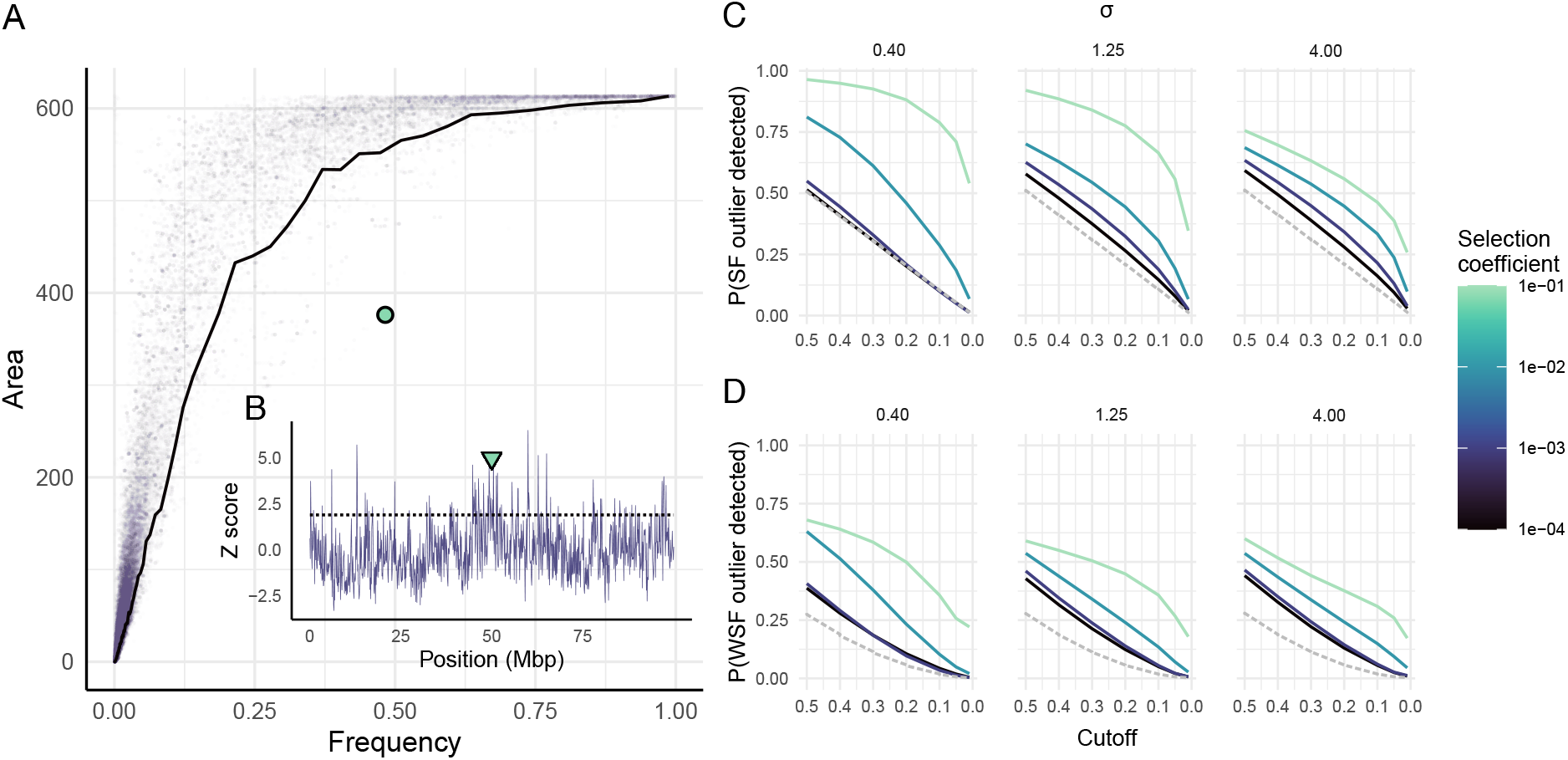
Power to detect selective sweeps using continuous space. (A) Example genomewide joint distribution between frequency and area of all variants 195 time steps after the introduction of an allele with a selection coefficient of *s* = 0.1 in a population with a mean per-generation dispersal distance of 0.40. The black line represents the tenth percentile cutoff for SF outliers; the sweeping allele is highlighted in green. (B) Example genome-wide windowed analysis of the ratio of SF outliers to non-outliers at the same time point in the same simulation. The dotted line represents the tenth percentile cutoff for WSF outliers; the green triangle highlights the locus of the sweeping allele. (C) Probability of detecting the sweeping allele as a SF outlier as the strictness of the outlier cutoff increases along the x axis. Panels represent *σ* values 0.40, 1.25, and 4.0; line color represents the strength of selection. The dashed gray line represents the proportion of the genome identified as SF outliers. (D) Probability of detecting the sweeping allele as a WSF outlier in the same simulations; the dashed gray line represents the proportion of the genome identified as WSF outliers.

We characterized power of the spatial-frequency (SF) outlier and windowed (WSF) outlier methods using forward-in-time spatial simulations for differing combinations of selection coefficients and dispersal rates (*σ*) while varying the stringency of the empirical cutoff used to define outliers. Unsurprisingly, we found that our method is best powered to detect the sweeping allele when the strength of selection is strong and dispersal is limited (Figs. 3C and 3D). Using an empirical cutoff at the 10^*th*^ percentile leads to a reasonable balance of power and false positive rate given the parameterization of our simulations (Figs. 3C and 3D), here observed as an inflection point in our power curves. We thus chose this cutoff for later empirical analysis.

Across all dispersal distances and selection coefficients, the probability of detecting the sweeping allele changes as a function of its frequency and increases sharply as the allele reaches fixation (Figure S2). Because there are fewer high-frequency SNPs overall and the landscape area they occupy is inherently high, we become more likely to identify the sweeping allele as an outlier in the joint distribution. Variation that is linked to this sweeping allele is also likely to take up less area relative to its frequency, contributing to power to detect the genomic window containing the sweeping allele. Interestingly, our method does not substantially suffer as a result of reduced sample size, and we observe only small reductions in power to detect a sweeping allele at low and high frequencies (Figure S3). At low frequencies, this is caused by the reduced probability of sampling allele carriers; at high frequencies, this is caused by the reduced probability of sampling non-carriers. The high power to detect positively-selected alleles at moderate frequencies even within small datasets may be attributed to both observed area and frequency being dependent on sampling effort, and thus their relationship to each other being relatively unaffected.

While our method demonstrates potential to detect in-progress sweeps, its empirical approach also results in a number of false positives that we control based on the defined outlier cutoff. For example, in our simulations with a single SNP under selection, identifying the lower tenth percentile of the joint area-frequency distribution as SF outliers and the upper tenth percentile of windowed SF outlier ratios as WSF outliers resulted in roughly 20, 000 SNPs (10% of genomic variation) being identified as SF outliers and roughly 3400 SNPs (less than 2% of genomic variation) being identified as WSF outliers. In empirical applications, genomic/biological annotations can help deal with sorting through these SNPs. Additionally, there are reasons beyond global positive selection that may cause SNPs to occupy a lower area relative to their frequency. For instance, local adaptation is likely to lead to this same pattern when examining populations across larger geographic regions. In empirical applications, disentangling the contributions of local and global adaptation to a variant’s position in the frequency-area distribution could be informed by examining the spatial distribution of geographic features alongside the distribution of the variant.

### Spatial signal identifies sweeping alleles in *Anopheles gambiae*

After demonstrating the potential of this spatial signal to identify positively selected variants, we next applied our approach to 687 West African *Anopheles gambiae* samples from the Ag1000G project (spatial sampling shown in Figure 4 inset; data described in The Anopheles gambiae 1000 Genomes Consortium et al. (2020)). Because this species is under strong selective pressures from human interventions, this is a useful system for testing our method with potential consequences for vector control efforts. For each SNP in the genome, we recorded its global frequency along with the locations (in latitude and longitude) of individuals carrying that SNP. After discarding SNPs that were found in fewer than three unique locations (706,991 alleles or approximately 71.2% of genomic variation, leaving 297,305 SNPs used in our analysis), we estimated the area on the landscape that each SNP occupies as the convex hull encompassing all individuals carrying the allele. Using the resulting distribution of SNP frequency and area, we used an empirical 10% area cutoff, conditioned on frequency, to identify SF outlier SNPs (Figure 4). We also describe each SF outlier’s distance from the mass of the joint frequency-area distribution using a *z* score calculated using similar-frequency variants (see Spatial genome scan).

**Figure 4.**
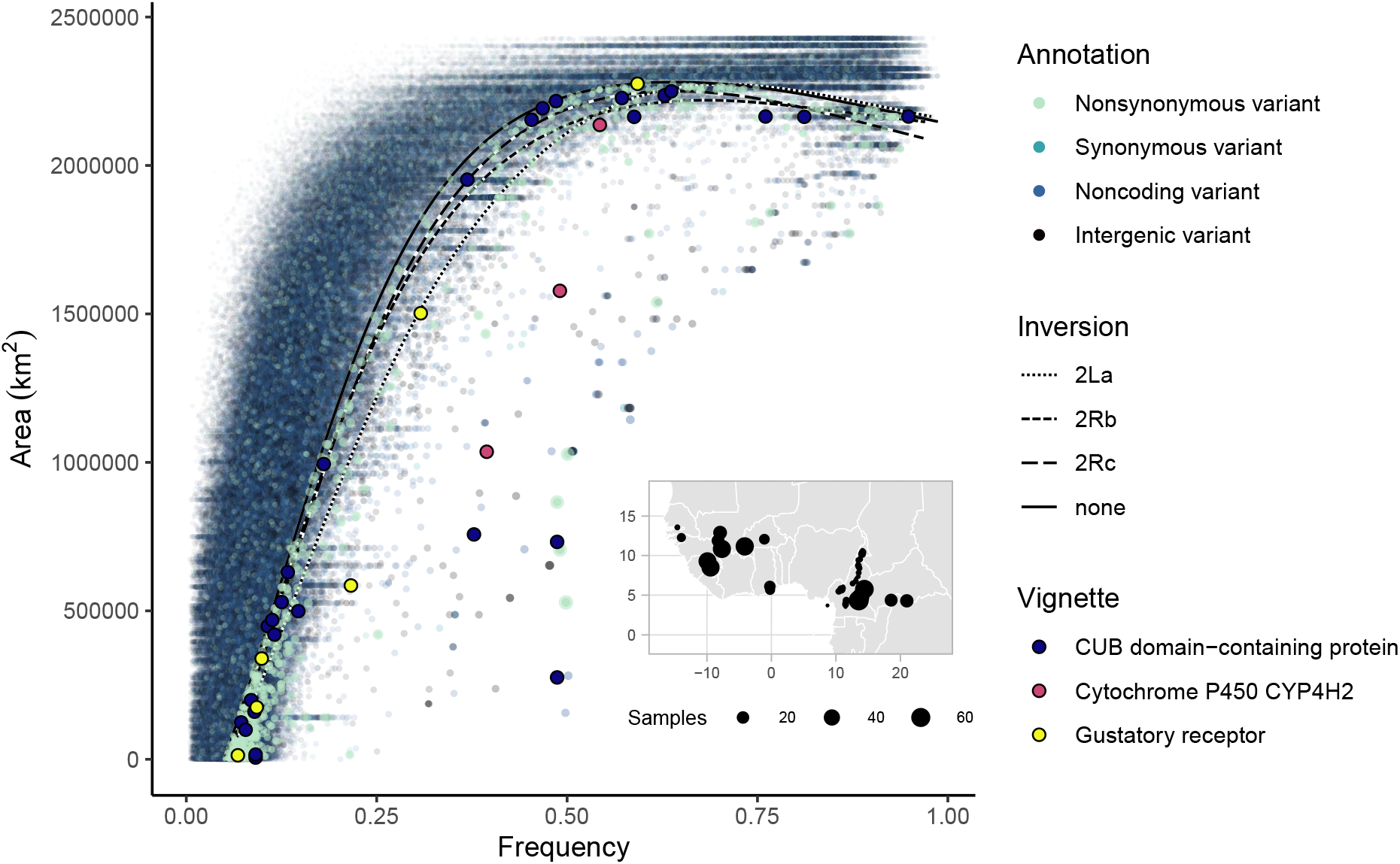
Joint distribution between frequency and area of all variants within the *Anopheles gambiae* genome. Each point represents a genomic variant; the point’s color indicates its annotation. Outlier variants highlighted in vignettes are colored and outlined. Black lines represent SF outlier cutoffs for each of the major inversions and the non-inverted portion of the genome, respectively. INSET: Sampling locations of *An. gambiae* samples. The size of the points represents the number of samples at each location.

As above, because positive selection reduces the area an allele occupies conditional on its frequency, we focused on SNPs in the joint distribution that match this pattern (Table S1). As linkage between SNPs might lead to genetic hitchiking between neutral and non-neutral SNPs, we further refined our analysis using a genomic scan to identify 100 kb windows of the genome that contain an excess of SF outliers after masking low-recombination regions, labeling SF outliers within these regions as windowed SF outliers (WSF outliers, Figure 5). To reduce the impact of other non-neutral processes on our analysis, we identified outliers separately for the major inversions (2La, 2Rb, 2Rc) and the rest of the genome, respectively, then SF and WSF outliers were pooled for downstream analyses. In total, we identified 36,500 SNPs as SF outliers and 3,875 SNPs as WSF outliers. It’s worth noting that our WSF outlier statistic agrees to some degree with previous searches for selective sweeps done in this system, and we find weak but significant correlations with windowed iHS statistics (*p* = 4.658 × 10^−8^; Anopheles gambiae 1000 Genomes Consortium (2017)), windows identified using partialSHI/C (*p* = 1.78 × 10^−16^; Xue et al. (2021)), and windows identified using PCADAPT (*p* = 1.07 × 10^−22^; Privé et al. (2020)), providing independent evidence that we are indeed picking up signals of positive selection.

**Figure 5.**
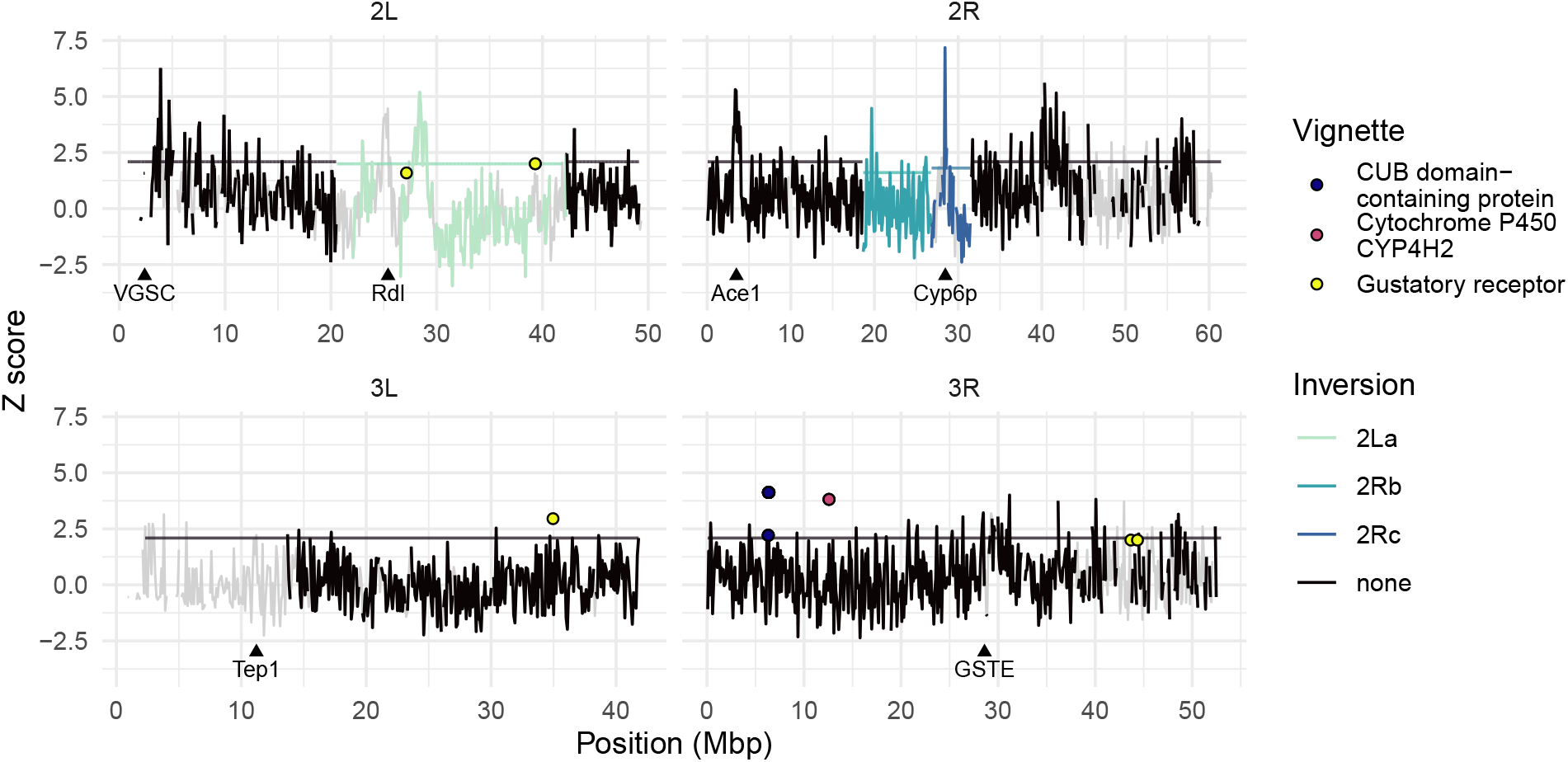
Genome-wide windowed analysis of the ratio of SF outliers to non-outliers. Flat horizontal lines represent WSF cutoffs for each of the major inversions and the non-inverted portion of the genome, respectively. Known loci associated with insecticide resistance and outlier variants highlighted in vignettes are labeled.

To assess what biological signals were being identified through our spatial approach, we first investigated the SF outliers’ annotations and associated genes. SF outliers were significantly (though modestly) enriched for coding SNPs (5.86% of SF outliers were located in coding regions compared to 5.31% of all SNPs retained in our analysis), with enrichments for both synonymous (*p* = 7.196 × 10^−7^, *χ*^2^ test) and nonsynonymous variants(*p* = 0.046, *χ*^2^ test) (Figure 4). Furthermore, SF outlier SNPs that are strong outliers in the joint genome-wide distribution of frequency and area fall within a number of loci associated with insecticide resistance, including an intronic variant within the *VGSC* locus (2L:2395567, *z* = −6.66; Martinez-Torres et al. (1998)), 31 intronic variants and 6 synonymous variants within the *Cyp6p* locus (2R:28480576-28505816, *z* scores range from −0.82 to −7.71; Müller et al. (2008)), and 29 intronic variants within the *Rdl* locus (2L:25363652 - 25434556, *z* scores range from −0.66 to −13.21 × 10^10^; Du et al. (2005)). Beyond these loci, the genes containing SF outliers (both coding and noncoding) were significantly enriched after Bonferroni correction for a number of GO biological processes important for sensory processing, including action potential (2.84 times enrichment, *p* = 1.64 × 10^−02^), neuron recognition (2.63 times enrichment, *p* = 1.92 × 10^−2^), and axon guidance (2.38 times enrichment, *p* = 1.21 × 10^−13^) (Table S2). Sensory perception and processing is often implicated in mechanisms of insecticide resistance in *Anopheles* species, and these genes are likely under further selective We next turned to our windowed WSF outliers (Figure 5). We again tested for enrichment of SNP annotations within these WSF outliers and found a significant enrichment of upstream (< 5kb from the transcription start site) gene variants (*p* = 8.004 × 10^−6^, *χ*^2^ test) and intronic variants (*p* = 1.015 × 10^−6^, *χ*^2^ test), which may indicate that a substantial portion of variants under positive selection are regulatory in this system. Further, ten WSF outliers fall within a well-known locus of insecticide resistance, Ace1 (2R:3484107-3495790, *z* scores range from −1.03 to −2.67). Genes containing WSF outliers were also significantly enriched for drug metabolism function (Enrichment score = 2.41, *p* = 0.0062) and enriched, though not significantly, for other relevant functions and annotations, including CRAL-TRIO domain-containing proteins, which are associated with eyesight (Enrichment score = 1.64, *p* = 0.47; Smith and Briscoe (2015)) and G protein-coupled receptors (Enrichment score = 1.78, *p* = 0.96, Bonferroni correction) (Table S3).

Finally, we searched for nonsynonymous outlier SNPs that could potentially be associated with functional changes. Investigating the functional consequences of nonsynonymous outlier SNPs, we found 37 WSF outliers and 237 SF outliers that impact protein stability (274 in total out of the 526 nonsynonymous outliers identified; Table S4). Among these are 44 variants within genes potentially associated with insecticide resistance as defined by the Ag1000g Selection Atlas, including methylenetetrahydrofolate reductase (AGAP007479:Y174D, *z* = −0.62), heme peroxidase 7 (AGAP004036:E564D, *z* = −0.86), and cytochrome P450 enzymes CYP6N2 (AGAP008206:K392N, *z* = −0.76), CYP4D15 (AGAP002418:L80M, *z* = −0.88), and CYP4H27 (AGAP008552, highlighted below) (Ag1000G Selection Atlas, 2017). The largest stability changes were observed within genes coding for metabolic proteins, which may be indicative of selective pressures exerted by insecticides, which target metabolic processes. High-magnitude stability changes are also observed within genes involved in sensory function, which may be due to changes in behavior associated with insecticide resistance (Thomsen et al., 2016), and genes involved in the regulation of gene expression. Changes in gene expression are often the mechanism behind insecticide resistance in *Anopheles* and other mosquitoes, and alterations in regulatory proteins may contribute to resistant phenotypes (Müller et al., 2008).

While there are a number of individual loci that we could highlight, in what follows we provide three particularly interesting vignettes of loci that represent strong spatial-frequency outliers in our scan and may be of functional consequence (see Figure 6).

**Figure 6.**
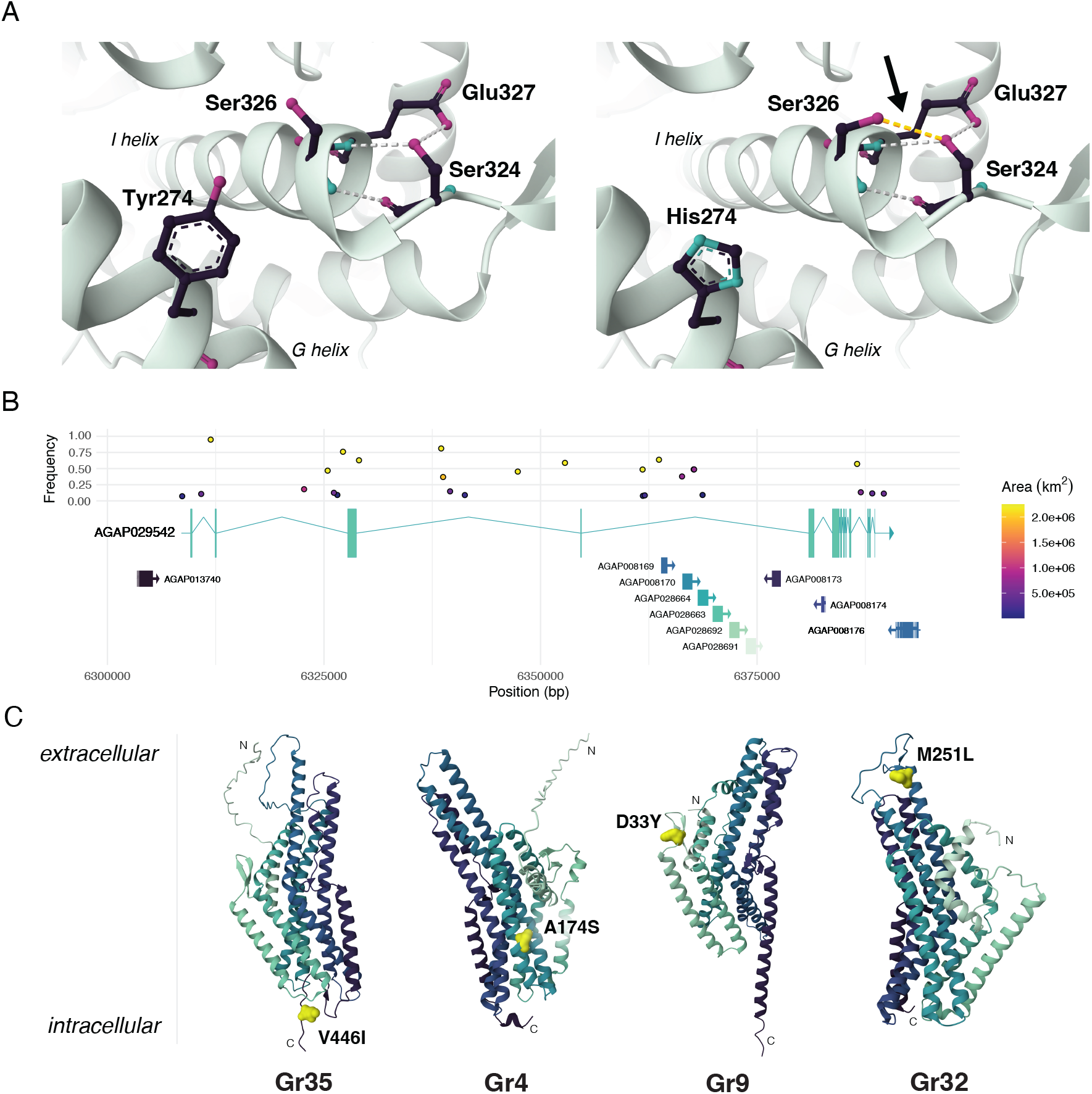
Impacts of outlier variants highlighted in vignettes. (A) Folding impacts of cytochrome P450 variant CYP4H27:Y274H. The left and right panels show position 274 and local structure for the ancestral tyrosine and the derived histidine, respectively. Relevant amino acids (S324, S326, E327) and hydrogen bonds are shown and the new hydrogen bond is highlighted in yellow. (B) Intronic outlier SNPs at the CUB domain-containing protein locus. Variants are shown at their genomic locations and colored by area; their y-axis position represents their frequency. Coding sequences and intronic segments for genes at the locus are annotated below; arrows represent the direction of transcription. (C) Structural locations of nonsynonymous variants in gustatory receptors. GRs are oriented with their extracellular domain upward and intracellular domain downward; chains are colored from light green at the N terminus to dark green at the C terminus and nonsynonymous mutations are highlighted in yellow.

### Cytochrome P450 *CYP4H27*

Cytochromes P450 are frequently implicated in the evolution of insecticide resistance across multiple taxa. In *Anopheline* mosquitoes, positive selection linked to Cyp450-associated resistance is most often associated with increased protein expression level due to copy number or transcriptional changes, rather than changes in protein function (Müller et al., 2008). However, using our SF outlier approach, we identify three nonsynonymous variants within one cytochrome P450 *CYP4H27* (Gene ID AGAP008552) that impact the folding stability and binding site conformation of this protein (Figure 6A). The first variant of interest, 3R:12573245T*>*C, is found only in five sampling locations, but is at a global frequency of almost 40%, and thus represents a highly significant outlier from the genome-wide joint distribution, the strength of which we can characterize with a *z*-score (*z* = −12.61, see Spatial genome scan). This mutation results in a shift from tyrosine to histidine at amino acid position 274 (Y274H) within the G helix of the CYP4H27 protein. The substitution shifts the orientation of the S326 residue on the N-terminal end of the I helix within the protein’s active site, allowing it to join a helix-stabilizing hydrogen bond between the residues of E327 and S324 and the carboxyl group of E27 and reducing the volume of the ligand binding pocket from 630.271 Å^3^ to 595. 784 Å^3^ as modeled by AlphaFold2 predictions (Ye et al., 2024, Abramson et al., 2024). Given that positively selected Cyp450 variants tend to be associated with increased protein expression that aids with insecticide detoxification, this modification to the structure of the CYP4H27 active site may have implications for its enzymatic activity, however further investigation is required to confirm this.

The other two variants of interest, 3R:12573798A*>*C and 3R:12573814T*>*A, are at 54% and 49% frequency (*z* = −2.62 and *z* = −10.36), respectively, and both result in charged amino acid changes between the L and M helices: 3R:12573798A*>*C causes a negatively charged serine at position 458 to change to a positively charged arginine (S458R), and 3R:12573814T*>*A causes a nonpolar valine at position 463 to change to a negatively charged aspartic acid (V463D). Among the individuals analyzed, we identified three mosquitoes with both the Y274H and S458R mutations segregating on the same haplotype and three mosquitoes with both Y274H and V463D segregating together, though we were unable to definitively find one haplotype with all three variants. AlphaFold predictions of the S458R and V463D carrying structures suggest these amino acid substitutions occur at the entrance of the ligand binding pocket, shifting its conformation, and that these changes are amplified when in combination with the Y274H mutation. Though the phenotypic effects of these three Cyp450 variants needs to be further investigated *in vivo*, their strong signals of positive selection may indicate another Cyp450-mediated mechanism of insecticide resistance.

### CUB domain-containing protein

A second interesting vignette from our analysis of outlier SNPs was AGAP029542, a relatively undescribed CUB domain-containing protein located on chromosome arm 3R (Figure 6B). While this gene’s function is unknown in *An. gambiae*, its ortholog in *Anopheles stephensi*, XP 035915664.1, has recently been implicated in contributing to insecticide resistance (Kumar et al., 2024). Our analysis identified 27 WSF outlier variants within this gene, exclusively in noncoding regions. These variants are some of the strongest outliers in the joint distribution between frequency and area (individual *z* scores range from −0.60 to −30.25 with 21 SNPs having *z* scores below −1.0; Figure 4), and we positively identify as many as 13 variants segregating within the same individual. While we cannot directly associate these variants with phenotypic changes, we speculate that positively selected variants in this gene might be associated with changes in transcription. Noncoding variants are commonly identified in selection scans in *Anopheles* and are understood to impact transcriptional regulation, a key feature in known mechanisms of insecticide resistance (Müller et al., 2008). Further, the functional domains of this protein have been shown to be highly conserved across *Anopheles* species (Kumar et al., 2024), making noncoding regions a more likely candidate for positive selection. Though more investigation is necessary to uncover the potential role of AGAP029542 in *An. gambiae* insecticide resistance, the density of outlier variants within AGAP029542 uncovered using our analysis highlights the strength of selection scans for generating hypotheses.

### Gustatory receptors

Chemosensory genes play a critical role in regulating insect behavior, and their evolution is implicated in broad phylogenetic as well as population-level adaptation such as avoidance or preference) of compounds such as those derived from plant secondary metabolites (e.g., Matsuo et al., 2007). Beyond identifying an enrichment of sensory genes amongst SF and WSF outliers, we identified six nonsynonymous mutations that were inferred to impact the stability of five gustatory receptors (Figure 6C). These proteins are cell-surface GPCRs, consisting of an extracellular N-terminus and ligand-interacting domain, seven transmembrane alpha helices, and an intracellular N-terminus and G protein-interacting domain. Two mutations we identified, Gr32:M251L (*z* = −1.79) and Gr9:D33Y (*z* = −0.98), are both located in the extracellular ligand-interacting domains of their respective proteins and result in nonconservative amino acid substitutions (shifting from sulfur-containing methionine to aliphatic leucine and acidic aspartate to aromatic tyrosine, respectively). Another outlier mutation within Gr32, Gr32:V195M (*z* = −2.63), is located proximate an extracellular ligand-interacting domain and results in a substitution from aliphatic valine to sulfur-containing methionine. These substitutions of amino acids with different biochemical properties in their extracellular domains may impact the receptor functions of these chemosensory proteins.

The other three nonsynonymous mutations we identified are more proximate to the intracellular domain, with two (Gr56:G147V, *z* = −0.76 and Gr4:A174S, *z* = −0.41) located on the intracellular ends of transbmembrane helices and one (Gr35:V446I, *z* = −1.17) located on the C terminal end of the protein. The mutations within Gr56 and Gr35 are both conservative substitutions of aliphatic amino acids to slightly larger ones and are predicted to cause small stability increases. Interestingly, the substitution of alanine with serine in Gr4 introduces a sulfur-containing amino acid to the intracellular region that destabilizes the protein, however this variant is observed at relatively high frequency (59.3%) and area (2.28e6 km^2^), indicating that this outlier variant is potentially under weaker selection than the others. While these gustatory receptors may be evolving in *An. gambiae* due to insecticide-mediated pressures, there are also broad biological reasons for chemosensory proteins to be evolving on a macroevolutionary scale within *Anopheline* mosquitoes.

## Discussion

Tests for positive selection are a common tool in population genetic analyses - however, the signals associated with positive selection, such as shifts in the site frequency spectrum and levels of observed heterozygosity, are often also impacted by spatial population structure (Battey et al., 2020a, Chotai et al., 2024). In this paper, we leverage such spatial structure to identify variants under positive selection using the joint distribution between allele frequency and area as our signal. Through simulation, we show how the increase in frequency associated with positive selection impacts an allele’s spread through space relative to its neutral counterparts, causing a striking shift in the expected frequency-area distribution. We then use this observation to develop a method for identifying sweeping alleles by identifying outliers from the genomic distribution of variants’ frequency and area. After demonstrating our method’s power to identify sweeping alleles in simulated datasets, we apply our approach to samples from the Ag1000G dataset to discover candidate variants undergoing positive selection. By considering spatial population structure, we uncover previously undescribed, epidemiologically relevant variants that display spatial signals of positive selection, demonstrating the potential for spatial analysis in uncovering evolutionary processes.

Our approach is based on the observation that, conditioning on frequency, variants that are positively selected take up less area on a continuous landscape than neutral variants. This is due to the fact that while positively selected alleles spread through a population more quickly than neutral ones, their spatial extent is constrained relative to their increasing frequency. In contrast, neutral alleles that are not lost can be expected to diffuse across the landscape while remaining at low frequency, creating a dramatic difference between the joint frequency-area trajectory of selected and neutral variants. This effect dissipates as selection and spatial structure get weaker: for the former, this is due to the sweeping allele being less likely to increase in frequency, and for the latter, the spatial spread of the sweeping allele is less restricted. While we modeled this frequency-area signal in simulations of a globally beneficial allele undergoing a selective sweep, there are other evolutionary phenomena that could lead to variants occupying a small area while being at high frequency. In particular, local adaptation would produce similar signals due to a beneficial allele being geographically restricted. The salient difference here is that an allele, rather than being globally beneficial across the landscape, may only be beneficial over a subset of the range, however from the standpoint of our frequency-area distribution calculated at a static point in time the two may be indistinguishable. This raises the possibility that some fraction of the candidate alleles which we have identified as outliers in the Ag1000G dataset may represent local adaptation, rather than in-progress global sweeps.

Our approach is in some sense analogous to earlier work which sought to contrast the frequency of an allele with its age. For instance, Slatkin and Bertorelle (2001) compared allele frequency to linked variation to identify the signature of selection, as neutral alleles at higher frequencies should be linked to relatively more polymorphism. Similarly, haplotype scan statistics aimed at detecting selection such as EHH (Sabeti et al., 2002) or iHS (Voight et al., 2006) use as signal haplotypes that are long relative to their frequency because recombination has had little time to break apart sequence surrounding beneficial alleles that are pushed to high frequency by selection. Here, rather than search for alleles that are young relative to their frequency, we are searching for alleles that are spatially restricted relative to their frequency. Our method is conceptually similar, but focused on spatial rather than temporal information.

At its most basic, the methodology we use here is a form of outlier detection, which comes with it a number of well described issues (e.g. Smiti (2020) or in the more narrow context of genetics (Teshima et al. (2006)). Our simulations show that our method is relatively well-powered; by focusing on the lower tenth percentile of variants in the joint frequency-area distribution (identified as SF outliers), we are able to successfully identify a sweeping allele up to 79% of the time in highly structured populations, and this power is not substantially impacted by reducing the number of samples analyzed. However we expect there to be a concomitant high rate of false positives – 10 percent of genomic variation is always identified as outliers. We further refined our spatial analysis by taking advantage of the fact that sweeping variants bring along linked neutral alleles. Using a window-based *z*-score to identify genomic regions with an excess of SF outliers, we identified the sweeping allele up to 35% of the time at a selection coefficient of *s* = 0.1 while reducing the number of false positives to roughly 1.7% of genomic variants. Unsurprisingly, in simulation both approaches were most successful at identifying a sweeping allele when both spatial structure and selection pressure were strong, as this is when an allele’s spread is most constrained relative to its increasing frequency. The windowed approach reduces the number of outliers identified, making downstream analysis simpler, but spatial outliers that are not within outlier windows may still be under positive selection. While we found that a tenth percentile cutoff was the most effective at minimizing both the false negative and false positive rates when we applied our method to simulated data, our approach is constrained by the trade-off between the two. This is however not unique to spatial scans for selection, and high false negative and false positive rates occur in empirical applications of traditional selection scans, even under relatively simple demographic models (Teshima et al., 2006). Additionally, while our simulations used uniform recombination maps, we found in our *An. gambiae* analysis that regions with low recombination are prone to inflated *z*-scores due to there being fewer variants overall identified in a window. We removed these segments from our windowed analysis before setting the WSF outlier cutoff, however many of these masked regions were still found to have relevant SF outlier SNPs.

Ultimately the strength of a method for finding selection rests in its application. We applied our approach to 687 samples of the malaria vector *Anopheles gambiae* from West Africa in order to identify variants under positive selection throughout this range (The Anopheles gambiae 1000 Genomes Consortium et al. (2020)). Anopheles mosquitoes in West Africa are known to be under strong selective pressure due to human intervention, and consequently genes that are insecticide targets, as well as genes involved in metabolism, behavior, and cuticle formation are often identified as undergoing or having undergone selective sweeps (e.g. Clarkson et al., 2021, Xue et al., 2021). Further, surveillance of currently sweeping variants and discovery of uncharacterized ones is crucial for maintaining successful vector control efforts for this critical public health threat. By considering spatial population structure, our method uncovers a number of epidemiologically relevant variants that display signals of positive selection and have not yet been noted by conventional selection scans.

The suite of *An. gambiae* genes containing SF and WSF outliers is enriched for genes involving sensory processing - a mechanism implicated in behavioral responses to insecticide pressures - and insecticide metabolism. Further, we identify outliers in a number of IR-associated loci, including *Rdl, Ace1*, and *Cyp6p*, and we highlight three outlier signals of particular interest. The first is three structure-altering nonsynonymous variants in the cytochrome P450 *CYP4H27*, a member of a protein family associated with insecticide resistance. Though we were unable to link all three variants to one haplotype, the variant displaying the strongest signal of positive selection, CYP4H27:Y274H, has a considerable impact on the shape and size of the protein’s binding pocket when modeled using Alphafold3 (Abramson et al., 2024). The second vignette of interest occurs in a CUB-domain containing protein that has 27 WSF outlier variants within it. While the protein is undescribed in *An. gambiae*, it is orthologus to an IR-associated protein in *An. stephensi* (Kumar et al., 2024). All of these variants are noncoding, but are strong outliers in the joint frequency-area distribution, underscoring the potential importance of noncoding variants to adaptation in this system. The last vignette we highlight focuses on six stability-altering nonsynonymous variants in gustatory receptors. These variants tend to be radical amino acid substitutions and are primarily located on either extracellular receptor domains or intracellular protein-binding domains of these receptors, indicating potential changes to both ligand binding and cell signaling. *Anopheles* GRs experience selection pressures beyond those caused by human vector control measures, and this signal demonstrates how our method can identify positive selection across a wide range of potential selective pressures and functional consequences. These findings generate new hypotheses for potential avenues of adaptation in *An. gambiae* and demonstrate the potential of this method in identifying otherwise-overlooked sweeping alleles.

A major consideration in benchmarking the power of our method when applied to real populations is how appropriate the simulation was. This spatial signal assumes a population where dispersal is continuous across the landscape - i.e., there are no major barriers to dispersal and every individual is equally likely to disperse in every direction. Further we assumed that selection pressure is homogeneous across the landscape as well. Heterogeneity of dispersal and selection may have major impacts on how an allele spreads across a landscape, and should be taken into consideration when applying our method to real populations. As mentioned earlier, other types of positive selection could generate this signal, such as local adaptation, which would result in a high-frequency allele occupying a geographically restricted area. Further, our power analysis assumes relatively uniform sampling coverage across the landscape, which is unrealistic for real populations. Though we recover signal of selection in the unevenly sampled *An. gambiae* dataset, consideration of the spatial distribution of samples is also necessary for empirical applications of this method, and further simulation-based analysis should include exploration of different spatial sampling schemes, as uneven sampling has been shown to bias other spatial-genetic analyses (Rehmann et al., 2024). That being said, our SNP-based approach and its robust power at sample sizes much lower than in the *An. gambiae* dataset make this type of analysis applicable to many spatial population genetic datasets. Finally, as previously discussed, our approach strikes a balance between minimizing false negatives and false positives, and is likely to both overlook and falsely identify alleles as being under positive selection (Teshima et al., 2006). For the latter case, genome annotation in empirical applications can help in identifying biologically relevant signals.

Here, we focus on the dynamics of beneficial alleles in continuous space; more generally, spatially informed genetic analyses have great potential to pick up on signals obscured by population structure in traditional analyses. While further inquiry is necessary into how the spatial impacts of other forms of selection, such as local adaptation, impact the frequency-area trajectory of an allele, we anticipate that data such as historical samples or ecologically relevant climatic information could help differentiate between different forms of adaptation. Generally we see great potential for integrating spatial information into genetic scans for selection; as the genetic history of a population unfolds in time and space, it is increasingly possible to ask questions about the geographic origins of genetic variants and to unravel the spatial history of adaptation.

## Materials and Methods

### Spatial simulation

To simulate selective sweeps in continuous space, we used a `SLiM v3` (Haller and Messer, 2019) model described in Battey et al. (2020a) and elaborated on in Chevy et al. (2024), in which positive selection impacts the mortality of individuals carrying the focal allele. Individuals were simulated across a 25 × 25 square continuous landscape with a target equilibrium density of *κ* = 5 individuals per unit square area. A simulation time step consists of three subsequent stages, all of which are controlled by a dispersal and interaction parameter *σ*. In our simulations, *σ* was varied between 0.40, 1.25, and 4.0 to achieve initial Wright neighborhood sizes between 10, 100, and 1000 (though these neighborhood sizes changed as the sweeping allele spread and population density increased).

In the first stage of a time step, individuals reproduce, choosing a mate from distance *x*_*i*_ with a probability proportional to exp 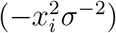. The number of offspring produced from a mating event is drawn from a Poisson distribution with a mean of 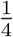. In the next stage, newly born individuals disperse from their focal parent at distances drawn from ∼ *N* (0, *σ*^2^). In the final stage, living individuals on the landscape compete locally for resources and space, dying with a probability *µ*(*u*) where *u* is the local population density. For an individual with neighbors (*x*_1_, *x*_2_, …*x*_*n*_), *u* is proportional to 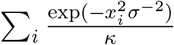, and at equilibrium population density, *µ*(*u*) is on average equal to the rate at which new individuals are born. Following Chevy et al. (2024), a positively-selected allele has additive effects such that an individual with *k* copies of the allele has a reduced mortality 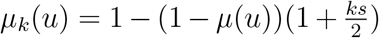. This form is chosen such that the positively-selected allele initially grows in copy number at a rate 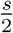. Because individuals carrying the allele have reduced mortality *µ*_*k*_(*u*),the spread of the positively-selected allele ultimately causes population density to increase (similar to what was observed in Chevy et al. (2024)).

Each individual has a diploid 10^8^ bp genome that recombines at a rate of 10^−8^ per bp per generation. After an initial burn-in of 1000 time steps, a positively selected mutation with a selection coefficient chosen from [1 × 10^−4^, 0.001, 0.01, 0.1] is added to the center of the genome of a newly born individual at the center of the landscape. As the simulation progresses, the frequency of the sweeping allele is tracked alongside the landscape area occupied by individuals carrying the sweeping allele, calculated as the area of the convex hull containing all allele carriers (Barber et al., 1996). We ran 100 total simulations to fixation across each parameter combination; simulations where the allele did not fix were restarted from the time point just before the addition of the sweeping allele.

### Power analysis

To assess the power of this spatial signal to identify variants under positive selection, we analyzed the spatial distribution of neutral variation across the genome over the course of our simulations. From each previously described simulation, we saved tree sequences over the course of the sweep (at a rate of 1 tree per 1 time step for a selection coefficient of 0.1, 2 time steps for 0.01, 5 time steps for 0.001, and 10 time steps for 1 × 10^−4^, respectively). Tree sequences were saved at different rates for different selection coefficients in order to capture the full dynamics of allele frequency while reducing computational time and memory load (Kelleher et al., 2018).

After simulating, we used msprime version 1.3.1 (Baumdicker et al., 2022) to recapitate the tree sequence, adding neutral mutations to the genome based on a population size equal to the population size at the tenth time step of the simulation (after which the population was spatially established), using a mutation rate of 10^−8^ per bp per generation. We then measured the frequency and area (through the same methods described in Methods Spatial simulation) of each variant across the genome, taking note of the sweeping allele. We used the joint distribution between allele frequency and occupied area of all variants across the genome to identify spatial-frequency (SF) outliers by binning every 1,000 variants (grouped by frequency), then classifying the lower *Nth* percentile in regards to area of these groups as spatial-frequency (SF) outliers (Figure 3 A). To assess our power to identify the sweeping allele based on the joint distribution between frequency and area, we tested for the presence of the sweeping allele among these SF outliers while varying the percentile cutoff between 0.5 and 0.01. We performed the same frequency-area analysis on tree sequences after down-sampling to 200 and 50 individuals, respectively, in order to assess our method’s power on smaller datasets.

Additionally, we analyzed the ratio of SF outliers to non-outliers in 100kb windows along the genome (Figure 3 B). To prevent our analysis from being thrown off by windows without many variants, we used as our statistic a *Z*-score defined as

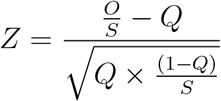

where *O* is the number of SF variants in a window, *S* is the total number of variants within a window, and *Q* is the genome-wide proportion of SF outliers to all variants. SF outlier SNPs within windows that had SF outlier:non-outlier ratios above the Nth percentile of genome-wide ratios were identified as windowed spatial-frequency (WSF) outliers while again varying the percentile cutoff between 0.5 and 0.01, and we again tested for whether the sweeping allele was identified as being within a WSF outlier window.

### *Anopheles gambiae* analysis

#### Data processing

We applied our method to whole genome sequencing of *Anopheles gambiae* available through the Ag1000G project (The Anopheles gambiae 1000 Genomes Consortium et al., 2020). Genotype data and metadata were downloaded from the Ag1000G phase 3 (Ag3.0) data release (accessed April 2024; https://malariagen.github.io/vector-data/ag3/ag3.0.html). VCFs relating to 28 sampling sets were merged using `bcftools` v1.20 (Danecek et al., 2021) and the command ‘merge’ with default options. Metadata were filtered to identify *An. gambiae* individuals as defined in the metadata header: ‘aim species’ and ‘taxon’. The merged VCF was filtered to retain only those *An. gambiae* individuals, resulting in 1470 individuals for downstream analyses. Monomorphic sites were removed using `bcftools` v1.20 and command ‘filter’ with options -e ‘AC==0 || AC==AN’. Finally, the VCF was filtered to retain high quality sites and accessible regions of the genome using the site filters available through Ag3.0. The final VCF contained 83,924,645 polymorphic sites, where 28% were classified as multiallelic (*>*1 alt allele).

The allelic state was polarized using the program `est-sfs` v2.03 (Keightley and Jackson, 2018) with *An. coluzzii, An. arabiensis, An. melas*, and *An. merus* as outgroup species. *An. coluzzii* and *An. arabiensis* are part of the Ag3.0 data release, and their VCFs were downloaded and processed in the same methods as outlined above. These VCFs were also filtered to retain only individuals with an unambiguous species identification. This left a total of 507 individuals of An. coluzzii and 368 individuals of *An. arabiensis*. The two other outgroup species, *An. melas* and *An. merus*, were downloaded from the available data sets of Fontaine et al. (2015). An input file for `est-sfs` was constructed using the custom script `estsfs_format.py` to partition the VCF into files with 100,000 sites for fast parallelization. The program est-sfs was then run with default parameters and the results integrated back into the *An. gambiae* VCF by adding an Ancestral Allele (AA) state to the INFO column of the VCF. The ancestral allele was determined to be the reference allele if the probability given by est-sfs was *>*0.90 and then the alternate allele if 1-probability was *>*0.90. Parsimony was used to polarize all other sites that could not be confidently polarized from the `est-sfs` results. Impacts of derived alleles were then annotated onto the VCF using SNPEff v5 and the AgamP4 reference genome (Assembly GCA 000005575.1) (Cingolani et al., 2012).

To reduce impacts of larger-scale population structure, we focused our analysis on 687 *An. gambiae* samples from West Africa grouped using `ADMIXTURE`. Ancestral clusters were estimated using `ADMIXTURE` (Alexander and Lange, 2011). A VCF of the genomic regions 3R:1-37,000,000 and 3L:15,000,000-41,000,000 (as noted in Anopheles gambiae 1000 Genomes Consortium (2017)) were filtered to retain SNPs with a minor allele frequency *>*5% and then thinned to retain only 1 site every 5,000bp. Thinning was done to reduce the influence of site dependency via linkage. The VCF was converted to bed format using `PLINK` v1.9 (Purcell et al., 2007). `ADMIXTURE` was run for a K from 1 to 5 (number of ancestral populations) with 5-fold cross-validation. Each `ADMIXTURE` analysis was repeated 30 times with different seeds. Final groups were visualized using `CLUMPAK` (Kopelman et al., 2015) and individuals were then assigned to one of the three clusters, the optimal K being 3 using an admixture proportion cutoff of 0.88.

A recombination map was estimated using `ReLERNN` v1.0.0 (Adrion et al., 2020) and mosquito sequences collected from the Central African Republic (CAR) (see Ag3.0 data release). The CAR population was used because sampling among the other *An. gambiae* populations was variable in both time and space. In contrast, CAR had 55 female mosquitoes collected during the same year (1994) and location (Bangui). `ReLERNN` was run on the phased and filtered VCF containing mosquitoes from CAR and a mask file (MASK) denoting the non-accessible regions of the genome. A custom demographic history (DEMO) trajectory for CAR mosquito population was inferred using `StairwayPlot2` (Liu and Fu, 2020). The classifier was trained using a window size of 26,000bp with default options and setting required parameters: –assumedMu 3.0*e* −9 –upperRhoThetaRatio 30 –phased −mask *MASK* – *n*DEMO. The resulting recombination maps were translated to cM/Mb and HapMap3 format using a custom script `rho2cMMb.py`.

### Spatial genome scan

We applied our analysis to 687 *An. gambiae* samples from West Africa grouped using `ADMIXTURE`. We recorded the frequency of each variant globally within the sample population alongside the area of the landscape occupied by its carriers. The area of the landscape occupied by allele carriers was determined by fitting a convex hull over the sampling locations of individuals carrying the allele through the same methods described in Methods section Spatial simulation, then calculating the area (in square kilometers) of the polygon determined by this hull using `pyproj.Geod`. Alleles that were not present in at least three unique locations were discarded from our analysis as we were unable to determine the area occupied by these variants.

From the joint distribution of allele frequency and area, we identified SF outliers as those below a spline fit (using the R package `npreg`) through the lower tenth area percentile of every 1000 SNPs binned by frequency after removing SNPs within the three major inversions (2La, 2Rb, 2Rc). To characterize the “distance” of an individual SNP from this cutoff, we used a modified *z* transformation of each SNP’s area compared to the distribution of areas within the defined 1000-SNP frequency bin, centering the areas according to the median of the distribution and then scaling by the median absolute deviation of the distribution. Such *z* scores give us a sense, although imperfect, of the strength of the outlier relationship. We then used a 100kb windowed analysis to identify WSF outliers beyond the 90th percentile of genomic *z* scores of SF outlier:non-outlier SNPs. This same approach was applied separately to SNPs within each the three major inversions (Figure S4). To prevent regions with low variation from inflating genome-wide *z* score values, we masked windows with a recombination rate below 1.5 cM/Mb before placing the 90th percentile cutoff to identify WSF outlier windows. We compared these windowed *z*-scores to the windowed values obtained from three other scans for selection: the windowed integrated haplotype score (iHS) (Ag1000G Selection Atlas, 2017), classification using partialSHI/C (Xue et al., 2021), and a windowed *z* score calculated using results from PCADAPT (Privé et al., 2020). We calculated iHS scores separately for each Ag1000G cohort included in our dataset, using windows calibrated to yield results where the 95% percentile of iHS values is at or below 0.1 (Malaria Genomic Epidemiology Network, 2022), then compared the average windowed iHS value across cohorts to our method’s windowed *z*-score value using a linear model. For partialSHI/C, we used windowed results from Xue et al. (2021), assigning windows classified as “Hard”, “HardPartial”, “Soft”, and “SoftPartial” a score of 1 and other windows a score of 0, then comparing these scores to our method using a linear model. Finally, we ran PCADAPT (using LD thinning and *K* = 10) on our dataset after removing the major inversions (2La, 2Rb, and 2Rc), which are in high linkage disequilibrium and would bias the results of this method (Privé et al., 2020). After correcting the *p*-values assigned to SNPs using the Bonferroni method, we identified outlier SNPs as those with a corrected *p*-value beneath 0.1, then calculated windowed *z*-scores using the same method described in Power analysis. We compared these windowed *z*-scores obtained from PCADAPT to those from our method using a linear model.

To assess potential phenotypic impacts of SF and WSF outliers, we searched for enrichment of SNP annotations obtained using SNPEff (Cingolani et al., 2012) using a *χ*^2^ test. Next, we identified genes containing SF and WSF outliers using VectorBase (Giraldo-Calderón et al., 2015), then used this list of genes to search for enrichment of functional annotations using either `PANTHER` (for SF outliers) or `DAVID` (for WSF and NS outliers) (Thomas et al., 2022, Mi et al., 2019, Dennis et al., 2003). Finally, for nonsynonymous WSF outliers, we obtained `.pdb` files of predicted protein structure for their associated genes using `AlphaFold3`, then used `ThermoMPNN` to assess how the associated amino acid substitution impacted protein stability (Jumper et al., 2021, Dieckhaus et al., 2024). For *CYP4H27*, we used the pairwise structure alignment tool available via www.rcsb.org/alignment with method `TM-align` to compare the ancestral and derived versions of the protein (Bittrich et al., 2024).

## Data availability

Code for simulations and analysis is available at the following link: https://github.com/kr-colab/spatialsweeps

## Acknowledgements

We thank Nate Pope, the members of the Kern-Ralph co-lab, Mara Lawniczak, and Matt Hahn for input and comments on the project and manuscript. CTR, STS, and ADK were funded by NIH Award R35GM148253 to ADK. PLR was funded in part by NIH Award R01HG012473.

## Supplementary material

### S1 Approximation to Fisher’s wave

In order to compare the allele frequency and area trajectories of our simulated sweeps to a deterministic model, we modeled an approximation of a positively-selected allele spreading across two-dimensional space as a traveling wave (Fisher, 1937, Kolmogorov et al., 1937). The positively-selected allele originates at the center of a 25×25-unit square*√*landscape and spreads in expanding circle whose radius *r* increases at a velocity 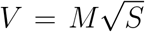 where *M* is the rate of per-generation dispersal (similar to the *σ* parameter in the SLiM simulation) and *S* is the selection coefficient of the allele. We can therefore define the radius*√*of the circle representing the extent of the sweeping allele at time point *t* as 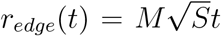 and use this to calculate the are*√*a occupied by the allele (Figure S1A). The width of the wavefront is defined as 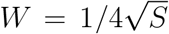 thus the spatial extent the fixed allele at time point *t* can be defined as 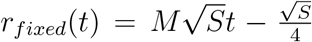. The circle representing the sweeping wavefront exits the landscape at *R* = 12.5 and completely encompasses the landscape at *R*_*max*_ *≈* 17.68. If *r*_*edge*_(*t*) *≤ R*, the area of the sweeping allele is simply *πr*_*edge*_(*t*)^2^. If *r*_*edge*_(*t*) *> R*, the area of the sweeping allele is defined as 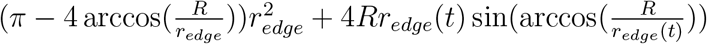.

To approximate the frequency of the allele, we modeled the sweeping wave in three dimensions as a conical frustum - a cone sliced horizontally - with height of 1 along the *z* axis, an upper radius *r*_*fixed*_, and base radius of *r*_*edge*_, and calculated its frequency out of the 25 × 25 × 1 volume of the landscape (Figure S1B). If *r*_*edge*_(*t*) < *R*, then the volume of the conical frustum, *V* (*t*), is equal to 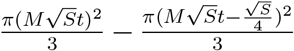. If *R ≤ r*_*edge*_ *≤ R*_*max*_ and the base begins to exit the landscape border, then

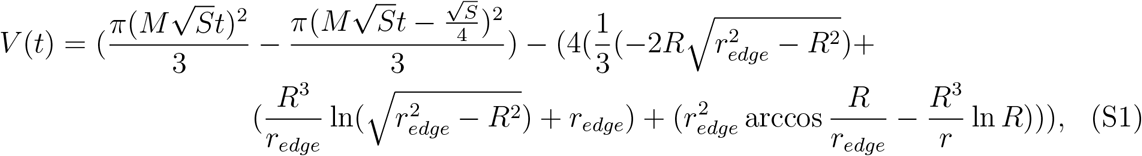

removing the volumes of the frustum that are outside the landscape. Finally, if *r*_*edge*_ *> R*_*max*_ and the base completely encompasses the landscape, the volume is calculated as the sum of the volume of a 25 × 25 rectangular prism with height 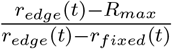and a conical frustum with height 1 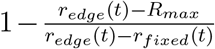, base radius *R*_*max*_, and upper radius *r*_*fixed*_ (*t*) (after subtracting off volumes of the frustum outside the landscape as described in Equation S1).

Our model corroborates both the logistic frequency trajectories of a sweep (Figure S1C) and the relationship between selection, frequency, and area that we observe in our continuous-space simulations (Figure S1D). We note that the key to the key to the nonlinear relationship between an allele’s frequency and area depends on the wavefront where the allele is still segregating; as the strength of selection increases and the width of the wavefront goes to zero the relationship between frequency and area becomes linear.

**Figure S1:**
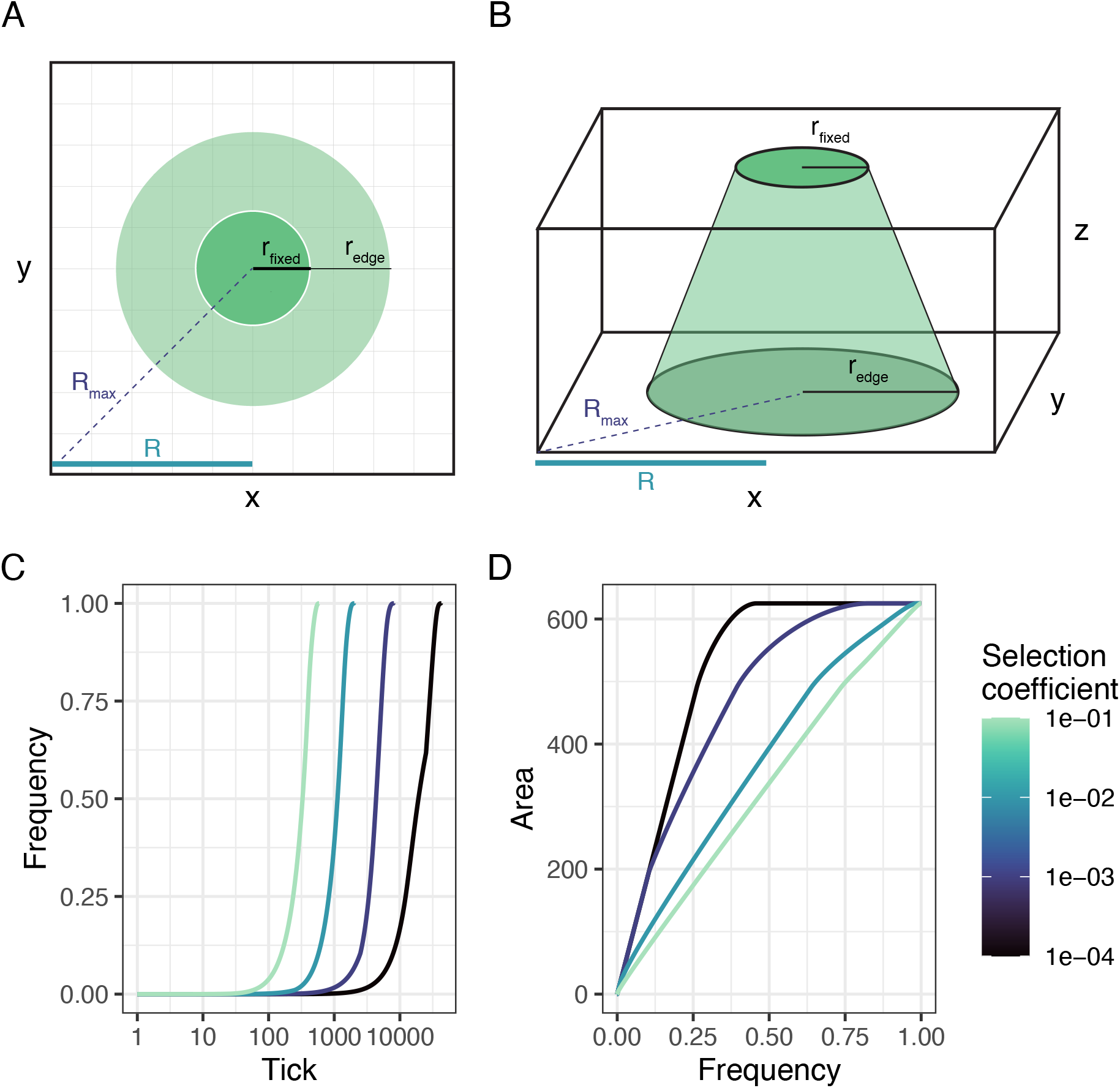
Approximation of a positively-selected allele spreading according to a Fisher wave across two-dimensional space. (A) Top-down view of the extent of the sweeping allele (*r*_*edge*_) and the extent of the allele’s fixation (*r*_*fixed*_). (B) Side view of the method used to approximate the frequency of the sweeping allele as the volume of the conical frustum defined by *r*_*edge*_ and *r*_*fixed*_ out of the volume of the landscape defined by *x, y*, and *z*. (C) Frequency of the positively-selected allele. (D) Joint distribution between frequency and area of the positively selected allele. In both plots, lines are colored by selection coefficient (*S*).

**Figure S2:**
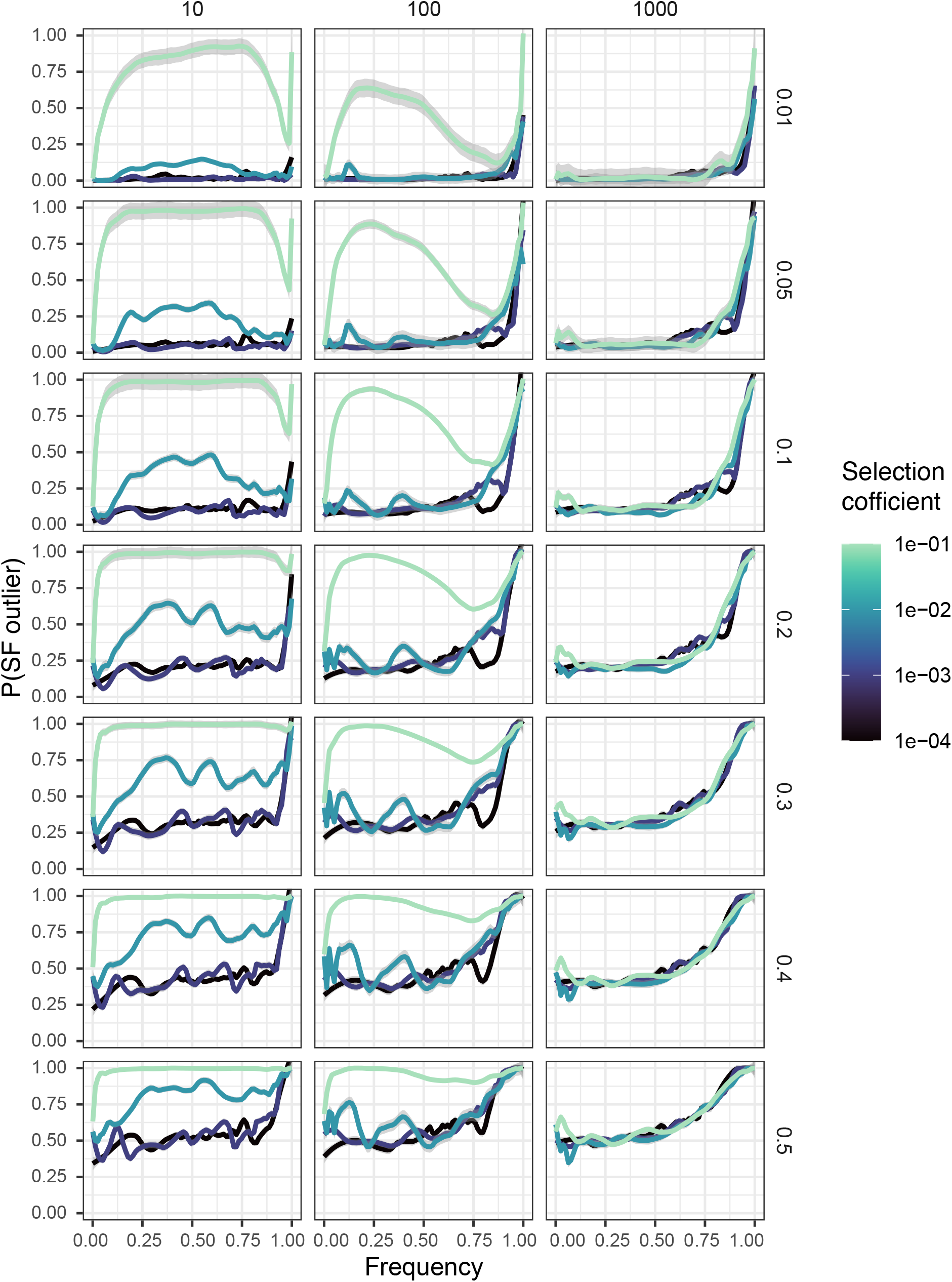
Probability of detecting a sweeping allele as a spatial outlier over the course of a sweep across dispersal distances (x axis) and cutoff percentiles (y axis). Lines are colored by selection coefficient. S3

**Figure S3:**
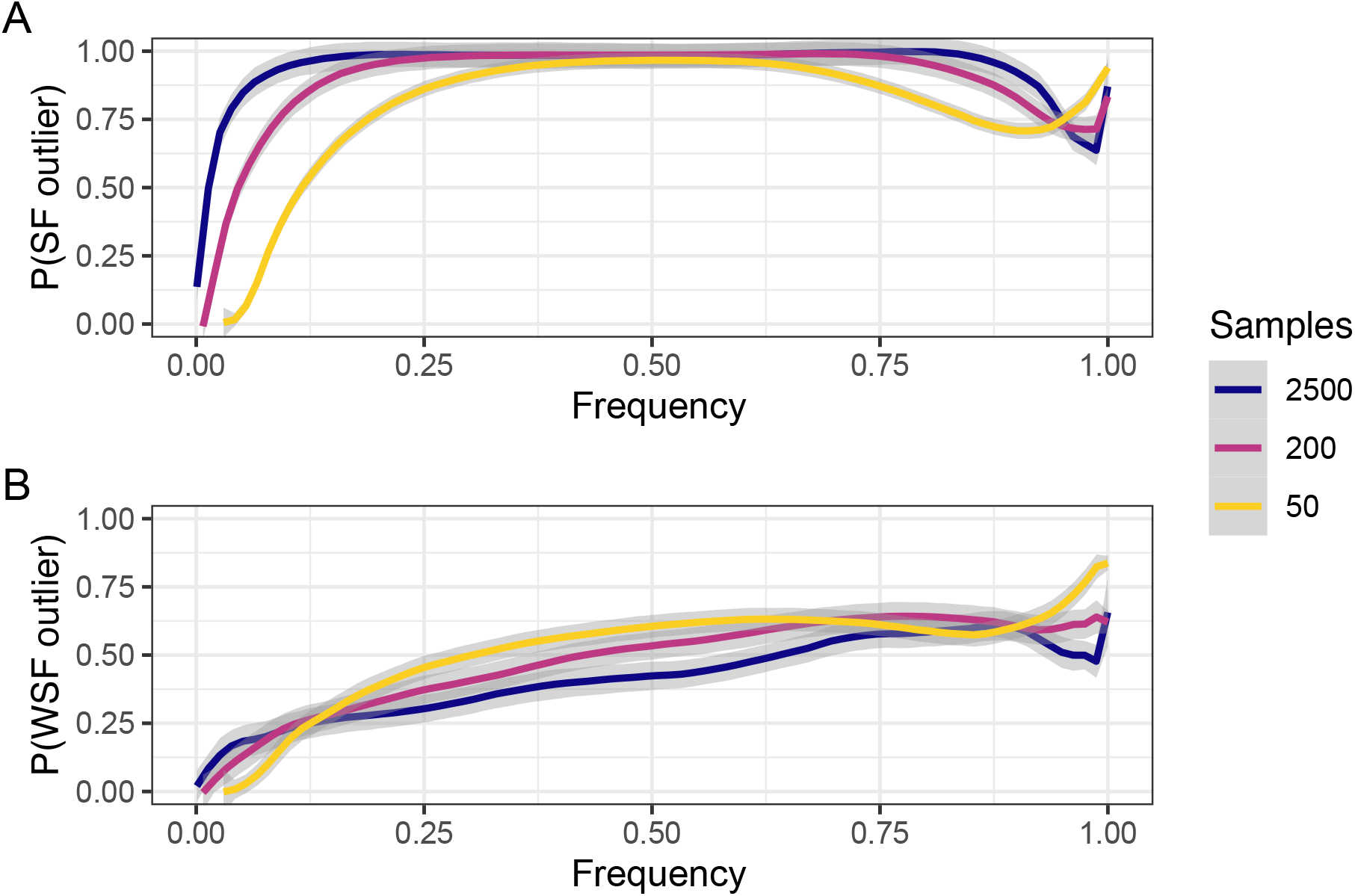
Probability of detecting a sweeping allele with selection coefficient *s* = 0.1 in a population with dispersal rate *σ* = 0.4 as a (A) spatial-frequency outlier and (B) WSF outlier over the course of a sweep at different sample sizes.

**Figure S4:**
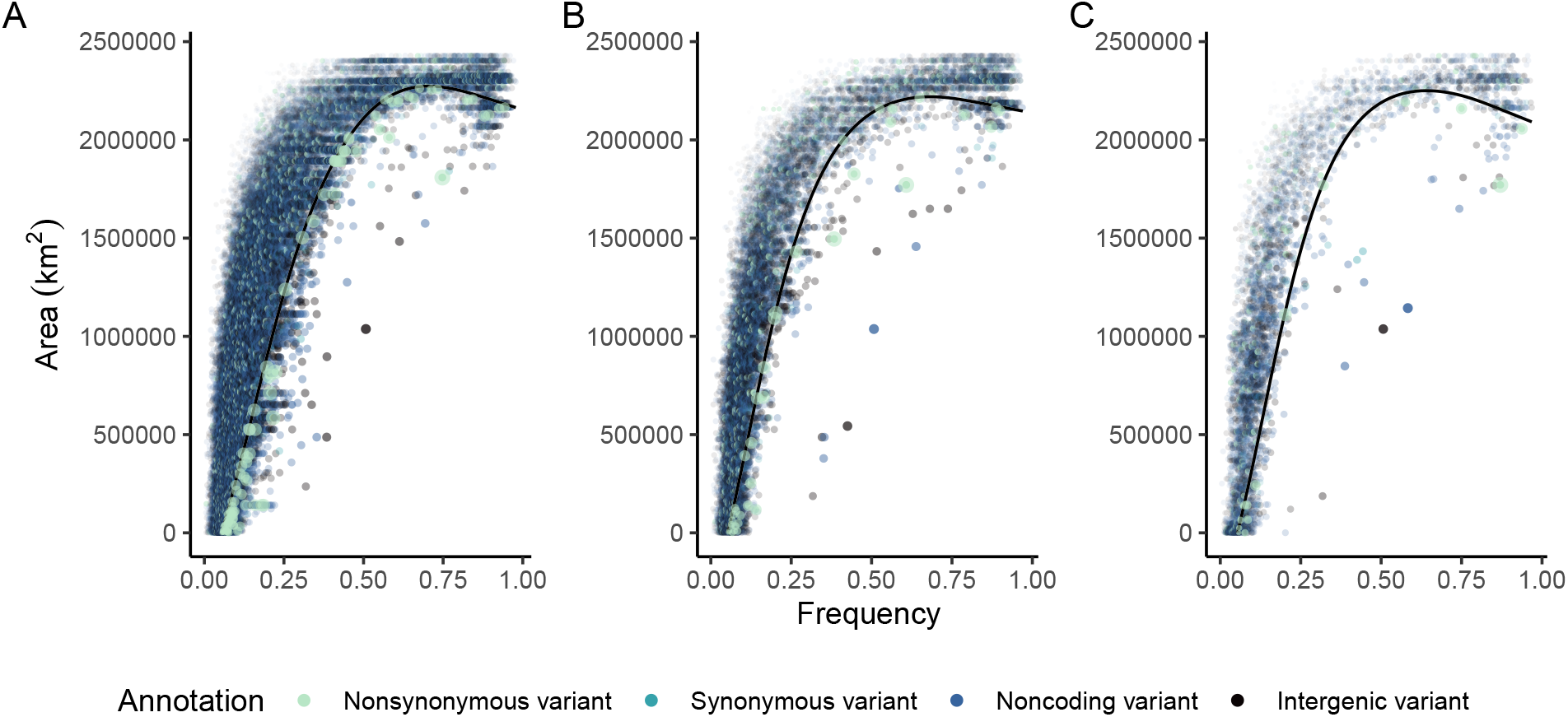
Joint frequency-area distribution for *An. gambiae* SNPs within the 2La (A), 2Rb (B), and 2Rc (C) inversions. SNPs are colored according to their SNPEff annotation; black lines represent the 10th percentile cutoff.

## References

[1] Josh Abramson, Jonas Adler, Jack Dunger, Richard Evans, Tim Green, Alexander Pritzel, Olaf Ronneberger, Lindsay Willmore, Andrew J. Ballard, Joshua Bambrick, and et al. Accurate structure prediction of biomolecular interactions with alphafold 3. Nature, 630(8016):493–500, May 2024. doi: 10.1038/s41586-024-07487-w.

[2] Jeffrey R Adrion, Jared G Galloway, and Andrew D Kern. Predicting the landscape of recombination using deep learning. Molecular biology and evolution, 37(6):1790–1808, 2020.

[3] Ag1000G Selection Atlas. Ag1000g selection atlas. https://malariagen.github.io/agam-selection-atlas/0.1-alpha3/, 2017. [Accessed 15-01-2025].

[4] Hussein Al-Asadi, Desislava Petkova, Matthew Stephens, and John Novembre. Estimating recent migration and population-size surfaces. PLoS genetics, 15(1):e1007908, 2019.

[5] David H Alexander and Kenneth Lange. Enhancements to the admixture algorithm for individual ancestry estimation. BMC bioinformatics, 12:1–6, 2011.

[6] Anopheles gambiae 1000 Genomes Consortium. Genetic diversity of the african malaria vector anopheles gambiae. Nature, 552(7683):96, 2017.

[7] Sandipan Paul Arnab, Md Ruhul Amin, and Michael DeGiorgio. Uncovering footprints of natural selection through spectral analysis of genomic summary statistics. Molecular Biology and Evolution, 40(7):msad157, 2023.

[8] C. Bradford Barber, David P. Dobkin, and Hannu Huhdanpaa. The quickhull algorithm for convex hulls. ACM Trans. Math. Softw., 22(4):469–483, December 1996. ISSN 0098-3500. doi: 10.1145/235815.235821. URL https://doi.org/10.1145/235815.235821.

[9] N. H. Barton, A. M. Etheridge, J. Kelleher, and A. Véber. Genetic hitchhiking in spatially extended populations. Theoretical Population Biology, 87:75–89, August 2013. ISSN 0040-5809. doi: 10.1016/j.tpb.2012.12.001. URL https://www.sciencedirect.com/science/article/pii/S0040580912001359.

[10] CJ Battey, Peter L Ralph, and Andrew D Kern. Space is the Place: Effects of Continuous Spatial Structure on Analysis of Population Genetic Data. Genetics, 215(1):193–214, May 2020a. ISSN 1943-2631. doi: 10.1534/genetics.120.303143. URL https://doi.org/10.1534/genetics.120.303143.

[11] CJ Battey, Peter L Ralph, and Andrew D Kern. Predicting geographic location from genetic variation with deep neural networks. eLife, 9:e54507, June 2020b. ISSN 2050-084X. doi: 10.7554/eLife.54507. URL https://doi.org/10.7554/eLife.54507. Publisher: eLife Sciences Publications, Ltd.

[12] Franz Baumdicker, Gertjan Bisschop, Daniel Goldstein, Graham Gower, Aaron P Ragsdale, Georgia Tsambos, Sha Zhu, Bjarki Eldon, E Castedo Ellerman, Jared G Galloway, Ariella L Gladstein, Gregor Gorjanc, Bing Guo, Ben Jeffery, Warren W Kretzschumar, Konrad Lohse, Michael Matschiner, Dominic Nelson, Nathaniel S Pope, Consuelo D Quinto-Cortés, Murillo F Rodrigues, Kumar Saunack, Thibaut Sellinger, Kevin Thornton, Hugo van Kemenade, Anthony W Wohns, Yan Wong, Simon Gravel, Andrew D Kern, Jere Koskela, Peter L Ralph, and Jerome Kelleher. Efficient ancestry and mutation simulation with msprime 1.0. Genetics, 220(3):iyab229, March 2022. ISSN 1943-2631. doi: 10.1093/genetics/iyab229. URL https://doi.org/10.1093/genetics/iyab229.

[13] Samir Bhatt, DJ Weiss, E Cameron, D Bisanzio, B Mappin, U Dalrymple, KE Battle, CL Moyes, A Henry, PA Eckhoff, et al. The effect of malaria control on plasmodium falciparum in africa between 2000 and 2015. Nature, 526(7572):207–211, 2015.

[14] Sebastian Bittrich, Joan Segura, Jose M Duarte, Stephen K Burley, and Yana Rose. Rcsb protein data bank: exploring protein 3d similarities via comprehensive structural alignments. Bioinformatics, 40(6): btae370, 06 2024. ISSN 1367-4811. doi: 10.1093/bioinformatics/btae370. URL https://doi.org/10.1093/bioinformatics/btae370.

[15] Hua Chen, Nick Patterson, and David Reich. Population differentiation as a test for selective sweeps. Genome research, 20(3):393–402, 2010.

[16] Elizabeth T. Chevy, Jiseon Min, Victoria Caudill, Samuel E. Champer, Benjamin C. Haller, Clara T. Rehmann, Chris C. R. Smith, Silas Tittes, Philipp W. Messer, Andrew D. Kern, Sohini Ramachandran, and Peter L. Ralph. Population genetics meets ecology: a guide to individual-based simulations in continuous landscapes, July 2024. URL https://www.biorxiv.org/content/10.1101/2024.07.24.604988v1. Pages: 2024.07.24.604988 Section: New Results.

[17] Meera Chotai, Xinzhu Wei, and Philipp W. Messer. Signatures of selective sweeps in continuous-space populations, July 2024. URL https://www.biorxiv.org/content/10.1101/2024.07.26.605365v1. Pages: 2024.07.26.605365 Section: New Results.

[18] Pablo Cingolani, Adrian Platts, Le Lily Wang, Melissa Coon, Tung Nguyen, Luan Wang, Susan J Land, Xiangyi Lu, and Douglas M Ruden. A program for annotating and predicting the effects of single nucleotide polymorphisms, snpeff: Snps in the genome of drosophila melanogaster strain w1118; iso-2; iso-3. fly, 6 (2):80–92, 2012.

[19] Richard M Clark, Eric Linton, Joachim Messing, and John F Doebley. Pattern of diversity in the genomic region near the maize domestication gene tb1. Proceedings of the National Academy of Sciences, 101(3): 700–707, 2004.

[20] Chris S Clarkson, Alistair Miles, Nicholas J Harding, Andrias O O’Reilly, David Weetman, Dominic Kwiatkowski, Martin J Donnelly, and Anopheles gambiae 1000 Genomes Consortium. The genetic architecture of target-site resistance to pyrethroid insecticides in the african malaria vectors anopheles gambiae and anopheles coluzzii. Molecular ecology, 30(21):5303–5317, 2021.

[21] Petr Danecek, James K Bonfield, Jennifer Liddle, John Marshall, Valeriu Ohan, Martin O Pollard, Andrew Whitwham, Thomas Keane, Shane A McCarthy, Robert M Davies, and Heng Li. Twelve years of SAMtools and BCFtools. GigaScience, 10(2), 02 2021. ISSN 2047-217X. doi: 10.1093/gigascience/giab008. URL https://doi.org/10.1093/gigascience/giab008.giab008.

[22] Michael DeGiorgio and Zachary A Szpiech. A spatially aware likelihood test to detect sweeps from haplotype distributions. PLoS genetics, 18(4):e1010134, 2022.

[23] Michael DeGiorgio, Christian D Huber, Melissa J Hubisz, Ines Hellmann, and Rasmus Nielsen. Sweepfinder2: increased sensitivity, robustness and flexibility. Bioinformatics, 32(12):1895–1897, 2016.

[24] Glynn Dennis, Brad T Sherman, Douglas A Hosack, Jun Yang, Wei Gao, H Clifford Lane, and Richard A Lempicki. David: database for annotation, visualization, and integrated discovery. Genome biology, 4:1–11, 2003.

[25] Henry Dieckhaus, Michael Brocidiacono, Nicholas Z Randolph, and Brian Kuhlman. Transfer learning to leverage larger datasets for improved prediction of protein stability changes. Proceedings of the National Academy of Sciences, 121(6):e2314853121, 2024.

[26] W Du, TS Awolola, P Howell, LL Koekemoer, BD Brooke, MQ Benedict, M Coetzee, and L Zheng. Independent mutations in the rdl locus confer dieldrin resistance to anopheles gambiae and an. arabiensis. Insect Mol. Biol., 14(2):179–183, April 2005.

[27] Justin C Fay and Chung-I Wu. Hitchhiking under positive darwinian selection. Genetics, 155(3):1405–1413, 2000.

[28] R. A. Fisher. The Wave of Advance of Advantageous Genes. Annals of Eugenics, 7(4):355–369, 1937. ISSN 2050-1439. doi: 10.1111/j.1469-1809.1937.tb02153.x. URL https://onlinelibrary.wiley.com/doi/abs/10.1111/j.1469-1809.1937.tb02153.x. eprint: https://onlinelibrary.wiley.com/doi/pdf/10.1111/j.1469-1809.1937.tb02153.x.

[29] Sir Fisher, Ronald Aylmer. The genetical theory of natural selection. Oxford, Clarendon Press, 1930, 1930. URL https://www.biodiversitylibrary.org/item/69976. https://www.biodiversitylibrary.org/bibliography/27468.

[30] Michael C Fontaine, James B Pease, Aaron Steele, Robert M Waterhouse, Daniel E Neafsey, Igor V Sharakhov, Xiaofang Jiang, Andrew B Hall, Flaminia Catteruccia, Evdoxia Kakani, et al. Extensive introgression in a malaria vector species complex revealed by phylogenomics. Science, 347(6217):1258524, 2015.

[31] Gloria I Giraldo-Calderón, Scott J Emrich, Robert M MacCallum, Gareth Maslen, Emmanuel Dialynas, Pantelis Topalis, Nicholas Ho, Sandra Gesing, VectorBase Consortium, Gregory Madey, et al. Vectorbase: an updated bioinformatics resource for invertebrate vectors and other organisms related with human diseases. Nucleic acids research, 43(D1):D707–D713, 2015.

[32] Benjamin C Haller and Philipp W Messer. SLiM 3: Forward Genetic Simulations Beyond the Wright–Fisher Model. Molecular Biology and Evolution, 36(3):632–637, March 2019. ISSN 0737-4038. doi: 10.1093/molbev/msy228. URL https://doi.org/10.1093/molbev/msy228.

[33] Janet Hemingway, Hilary Ranson, Alan Magill, Jan Kolaczinski, Christen Fornadel, John Gimnig, Maureen Coetzee, Frederic Simard, Dabiré K Roch, Clément Kerah Hinzoumbe, et al. Averting a malaria disaster: will insecticide resistance derail malaria control? The Lancet, 387(10029):1785–1788, 2016.

[34] Joachim Hermisson and Pleuni S Pennings. Soft sweeps: molecular population genetics of adaptation from standing genetic variation. Genetics, 169(4):2335–2352, 2005.

[35] John Jumper, Richard Evans, Alexander Pritzel, Tim Green, Michael Figurnov, Olaf Ronneberger, Kathryn Tunyasuvunakool, Russ Bates, Augustin Žídek, Anna Potapenko, et al. Highly accurate protein structure prediction with alphafold. nature, 596(7873):583–589, 2021.

[36] Peter D Keightley and Benjamin C Jackson. Inferring the probability of the derived vs. the ancestral allelic state at a polymorphic site. Genetics, 209(3):897–906, 2018.

[37] Jerome Kelleher, Kevin R. Thornton, Jaime Ashander, and Peter L. Ralph. Efficient pedigree recording for fast population genetics simulation. PLOS Computational Biology, 14(11):1–21, 11 2018. doi: 10.1371/journal.pcbi.1006581. URL https://doi.org/10.1371/journal.pcbi.1006581.

[38] Andrew D Kern and Daniel R Schrider. diploS/HIC: An Updated Approach to Classifying Selective Sweeps. G3 Genes |Genomes| Genetics, 8(6):1959–1970, June 2018. ISSN 2160-1836. doi: 10.1534/g3.118.200262. URL https://doi.org/10.1534/g3.118.200262.

[39] Ken-Ichi Kojima and Therese M Kelleher. Survival of mutant genes. The American Naturalist, 96(891):329–346, 1962.

[40] Ken-Ichi Kojima and Henry E Schaffer. Survival process of linked mutant genes. Evolution, pages 518–531, 1967.

[41] A. Kolmogorov, I. Petrovskii, and N. Piscunov. A study of the equation of diffusion with increase in the quantity of matter, and its application to a biological problem. Byul. Moskovskogo Gos. Univ., 1(6):1–25, 1937. URL http://books.google.com/books?id=ikN59GkYJKIC&lpg=PP1&dq=A.N.%20Kolmogorov%3A%20Selected%20Works&client=firefox-a&pg=PA242#v=onepage&q=&f=false.

[42] Naama M Kopelman, Jonathan Mayzel, Mattias Jakobsson, Noah A Rosenberg, and Itay Mayrose. Clumpak: a program for identifying clustering modes and packaging population structure inferences across k. Molecular ecology resources, 15(5):1179–1191, 2015.

[43] Jatin Kumar, Ankit Kumar, Yash Gupta, Kapil Vashisht, Shivam Kumar, Arvind Sharma, Raj Kumar, Ashoke Sharon, Praveen K. Tripathi, Ram Das, Om Prakash Singh, Shailja Singh, Soumyananda Chakraborti, Sujatha Sunil, and Kailash C. Pandey. A cub and sushi domain-containing protein with esterase-like activity confers insecticide resistance in the indian malaria vector ¡em¿anopheles stephensi¡/em¿. Journal of Biological Chemistry, 300(10), Oct 2024. ISSN 0021-9258. doi: 10.1016/j.jbc.2024.107759. URL https://doi.org/10.1016/j.jbc.2024.107759.

[44] M Elise Lauterbur, Kasper Munch, and David Enard. Versatile detection of diverse selective sweeps with flex-sweep. Molecular Biology and Evolution, 40(6):msad139, 2023.

[45] Xiaoming Liu and Yun-Xin Fu. Stairway plot 2: demographic history inference with folded snp frequency spectra. Genome biology, 21(1):280, 2020.

[46] Malaria Genomic Epidemiology Network. Workshop 6 - detecting genes under recent positive selection. In Training course in data analysis for genomic surveillance of African malaria vectors. 2022. URL https://anopheles-genomic-surveillance.github.io/home.html.

[47] Joseph Marcus, Wooseok Ha, Rina Foygel Barber, and John Novembre. Fast and flexible estimation of effective migration surfaces. Elife, 10:e61927, 2021.

[48] D Martinez-Torres, F Chandre, MS Williamson, F Darriet, JB Bergé, AL Devonshire, P Guillet, N Pasteur, and D Pauron. Molecular characterization of pyrethroid knockdown resistance (kdr) in the major malaria vector anopheles gambiae s.s. Insect Mol. Biol., 7(2):179–184, May 1998.

[49] Takashi Matsuo, Shigeru Sugaya, Jyunichiro Yasukawa, Toshiro Aigaki, and Yoshiaki Fuyama. Odorantbinding proteins obp57d and obp57e affect taste perception and host-plant preference in drosophila sechellia. PLoS biology, 5(5):e118, 2007.

[50] John Maynard Smith and John Haigh. The hitch-hiking effect of a favourable gene. Genet. Res., 23 (1):23–35, February 1974. ISSN 0016-6723, 1469-5073. doi: 10.1017/S0016672300014634. URL https://www.cambridge.org/core/product/identifier/S0016672300014634/type/journalarticle.

[51] Huaiyu Mi, Anushya Muruganujan, Xiaosong Huang, Dustin Ebert, Caitlin Mills, Xinyu Guo, and Paul D Thomas. Protocol update for large-scale genome and gene function analysis with the PANTHER classification system (v.14.0). Nat. Protoc., 14(3):703–721, March 2019.

[52] Jiseon Min, Misha Gupta, Michael M Desai, and Daniel B Weissman. Spatial structure alters the site frequency spectrum produced by hitchhiking. Genetics, 222(3):iyac139, November 2022. ISSN 1943-2631. doi: 10.1093/genetics/iyac139. URL https://doi.org/10.1093/genetics/iyac139.

[53] Pie Müller, Emma Warr, Bradley J Stevenson, Patricia M Pignatelli, John C Morgan, Andrew Steven, Alexander E Yawson, Sara N Mitchell, Hilary Ranson, Janet Hemingway, Mark J I Paine, and Martin J Donnelly. Field-caught permethrin-resistant anopheles gambiae overexpress CYP6P3, a P450 that metabolises pyrethroids. PLoS Genet., 4(11):e1000286, November 2008.

[54] Rasmus Nielsen, Scott Williamson, Yuseob Kim, Melissa J. Hubisz, Andrew G. Clark, and Carlos Bustamante. Genomic scans for selective sweeps using SNP data. Genome Res., 15(11):1566–1575, November 2005. ISSN 1088-9051, 1549-5469. doi: 10.1101/gr.4252305. URL https://genome.cshlp.org/content/15/11/1566. Company: Cold Spring Harbor Laboratory Press Distributor: Cold Spring Harbor Laboratory Press Institution: Cold Spring Harbor Laboratory Press Label: Cold Spring Harbor Laboratory Press Publisher: Cold Spring Harbor Lab.

[55] Florian Privé, Keurcien Luu, Bjarni J Vilhjálmsson, and Michael G B Blum. Performing highly efficient genome scans for local adaptation with r package pcadapt version 4. Molecular Biology and Evolution, 37 (7):2153–2154, 04 2020. ISSN 0737-4038. doi: 10.1093/molbev/msaa053. URL https://doi.org/10.1093/molbev/msaa053.

[56] Shaun Purcell, Benjamin Neale, Kathe Todd-Brown, Lori Thomas, Manuel AR Ferreira, David Bender, Julian Maller, Pamela Sklar, Paul IW De Bakker, Mark J Daly, et al. Plink: a tool set for whole-genome association and population-based linkage analyses. The American journal of human genetics, 81 (3):559–575, 2007.

[57] Clara T Rehmann, Peter L Ralph, and Andrew D Kern. Evaluating evidence for co-geography in the anopheles–plasmodium host–parasite system. G3: Genes, Genomes, Genetics, 14(3):jkae008, 2024.

[58] Harald Ringbauer, Graham Coop, and Nicholas H Barton. Inferring recent demography from isolation by distance of long shared sequence blocks. Genetics, 205(3):1335–1351, 2017.

[59] Francois Rousset. Genetic differentiation and estimation of gene flow from F-statistics under isolation by distance. Genetics, 145(4):1219–1228, 1997.

[60] Pardis C Sabeti, David E Reich, John M Higgins, Haninah ZP Levine, Daniel J Richter, Stephen F Schaffner, Stacey B Gabriel, Jill V Platko, Nick J Patterson, Gavin J McDonald, et al. Detecting recent positive selection in the human genome from haplotype structure. Nature, 419(6909):832–837, 2002.

[61] Todd A Schlenke and David J Begun. Strong selective sweep associated with a transposon insertion in drosophila simulans. Proceedings of the National Academy of Sciences, 101(6):1626–1631, 2004.

[62] Daniel R Schrider and Andrew D Kern. S/hic: robust identification of soft and hard sweeps using machine learning. PLoS genetics, 12(3):e1005928, 2016.

[63] Montgomery Slatkin and Giorgio Bertorelle. The use of intraallelic variability for testing neutrality and estimating population growth rate. Genetics, 158(2):865–874, 2001.

[64] Chris CR Smith and Andrew D Kern. dispersenn2: a neural network for estimating dispersal distance from georeferenced polymorphism data. BMC bioinformatics, 24(1):385, 2023.

[65] Chris CR Smith, Silas Tittes, Peter L Ralph, and Andrew D Kern. Dispersal inference from population genetic variation using a convolutional neural network. Genetics, 224(2):iyad068, 2023.

[66] Chris CR Smith, Gilia Patterson, Peter L Ralph, and Andrew D Kern. Estimation of spatial demographic maps from polymorphism data using a neural network. bioRxiv, 2024.

[67] Gilbert Smith and Adriana D. Briscoe. Molecular evolution and expression of the cral trio protein family in insects. Insect Biochemistry and Molecular Biology, 62:168–173, 2015. ISSN 0965-1748. doi: 10.1016/j.ibmb.2015.02.003. URL https://www.sciencedirect.com/science/article/pii/S0965174815000260.

[68] Abir Smiti. A critical overview of outlier detection methods. Computer Science Review, 38:100306, 2020.

[69] Lauren Alpert Sugden, Elizabeth G Atkinson, Annie P Fischer, Stephen Rong, Brenna M Henn, and Sohini Ramachandran. Localization of adaptive variants in human genomes using averaged one-dependence estimation. Nature communications, 9(1):703, 2018.

[70] Fumio Tajima. Statistical method for testing the neutral mutation hypothesis by dna polymorphism. Genetics, 123(3):585–595, 1989.

[71] Kosuke M Teshima, Graham Coop, and Molly Przeworski. How reliable are empirical genomic scans for selective sweeps? Genome research, 16(6):702–712, 2006.

[72] The Anopheles gambiae 1000 Genomes Consortium, Chris S. Clarkson, Alistair Miles, Nicholas J. Harding, Eric R. Lucas, C. J. Battey, Jorge Edouardo Amaya-Romero, Jorge Cano, Abdoulaye Diabate, Edi Constant, Davis C. Nwakanma, Musa Jawara, John Essandoh, Joao Dinis, Gilbert Le Goff, Vincent Robert, Arlete D. Troco, Carlo Costantini, Kyanne R. Rohatgi, Nohal Elissa, Boubacar Coulibaly, Janet Midega, Charles Mbogo, Henry D. Mawejje, Jim Stalker, Kirk A. Rockett, Eleanor Drury, Daniel Mead, Anna E. Jeffreys, Christina Hubbart, Kate Rowlands, Alison T. Isaacs, Dushyanth Jyothi, Cinzia Malangone, Maryam Kamali, Christa Henrichs, Victoria Simpson, Diego Ayala, Nora J. Besansky, Austin Burt, Beniamino Caputo, Alessandra della Torre, Michael Fontaine, H. Charles J. Godfray, Matthew W. Hahn, Andrew D. Kern, Mara K. N. Lawniczak, Samantha O’Loughlin, Joao Pinto, Michelle M. Riehle, Igor Sharakhov, Daniel R. Schrider, Kenneth D. Vernick, Bradley J. White, Martin J. Donnelly, and Dominic P. Kwiatkowski. Genome variation and population structure among 1,142 mosquitoes of the African malaria vector species Anopheles gambiae and Anopheles coluzzii, February 2020. URL https://www.biorxiv.org/content/10.1101/864314v2. Pages: 864314 Section: New Results.

[73] Paul D. Thomas, Dustin Ebert, Anushya Muruganujan, Tremayne Mushayahama, Laurent-Philippe Albou, and Huaiyu Mi. Panther: Making genome-scale phylogenetics accessible to all. Protein Science, 31(1):8– 22, 2022. doi: 10.1002/pro.4218. URL https://onlinelibrary.wiley.com/doi/abs/10.1002/pro.4218.

[74] Edward K Thomsen, Gussy Koimbu, Justin Pulford, Sharon Jamea-Maiasa, Yangta Ura, John B Keven, Peter M Siba, Ivo Mueller, Manuel W Hetzel, and Lisa J Reimer. Mosquito behaviour change after distribution of bednets results in decreased protection against malaria exposure. J. Infect. Dis., page jiw615, December 2016.

[75] Benjamin F Voight, Sridhar Kudaravalli, Xiaoquan Wen, and Jonathan K Pritchard. A map of recent positive selection in the human genome. PLoS biology, 4(3):e72, 2006.

[76] Scott H Williamson, Melissa J Hubisz, Andrew G Clark, Bret A Payseur, Carlos D Bustamante, and Rasmus Nielsen. Localizing recent adaptive evolution in the human genome. PLoS genetics, 3(6):e90, 2007.

[77] Sewall Wright. ISOLATION BY DISTANCE. Genetics, 28(2):114–138, March 1943. ISSN 1943-2631. doi: 10.1093/genetics/28.2.114. URL https://doi.org/10.1093/genetics/28.2.114.

[78] Alexander T Xue, Daniel R Schrider, and Andrew D Kern. Discovery of ongoing selective sweeps within anopheles mosquito populations using deep learning. Molecular biology and evolution, 38(3):1168–1183, 2021.

[79] Bowei Ye, Wei Tian, Boshen Wang, and Jie Liang. Castpfold: Computed atlas of surface topography of the universe of protein folds. Nucleic Acids Research, 52(W1):W194–W199, 05 2024. ISSN 0305-1048. doi: 10.1093/nar/gkae415. URL https://doi.org/10.1093/nar/gkae415.

